# Comparison of calling pipelines for whole genome sequencing: an empirical study demonstrating the importance of mapping and alignment

**DOI:** 10.1101/2022.09.18.508404

**Authors:** Raphael O. Betschart, Alexandre Thiéry, Domingo Aguilera-Garcia, Martin Zoche, Holger Moch, Raphael Twerenbold, Tanja Zeller, Stefan Blankenberg, Andreas Ziegler

## Abstract

Rapid advances in high-throughput DNA sequencing technologies have enabled the conduct of whole genome sequencing (WGS) studies, and several bioinformatics pipelines have become available. The aim of this study was the comparison of 6 WGS data pre-processing pipelines, involving two mapping and alignment approaches (GATK utilizing BWA-MEM2 2.2.1, and DRAGEN 3.8.4) and three variant calling pipelines (GATK 4.2.4.1, DRAGEN 3.8.4 and DeepVariant 1.1.0). We sequenced one genome in a bottle (GIAB) sample 70 times in different runs, and one GIAB trio in triplicate. The truth set of the GIABs was used for comparison, and performance was assessed by computation time, F_1_ score, precision, and recall. In the mapping and alignment step, the DRAGEN pipeline was faster than the GATK with BWA-MEM2 pipeline. DRAGEN showed systematically higher F_1_ score, precision, and recall values than GATK for single nucleotide variations (SNVs) and Indels in simple-to-map, complex-to-map, coding and non-coding regions. In the variant calling step, DRAGEN was fastest. In terms of accuracy, DRAGEN and DeepVariant performed similarly and both superior to GATK, with slight advantages for DRAGEN for Indels and for DeepVariant for SNVs. The DRAGEN pipeline showed the lowest Mendelian inheritance error fraction for the GIAB trios. Mapping and alignment played a key role in variant calling of WGS, with the DRAGEN outperforming GATK.

## Introduction

Whole genome sequencing (WGS) of a human genome for less than 1000 USD has become reality^1^, and cost might drop to even 100 USD with the Illumina NovaSeq system^2^. Large-scale WGS studies have already been initiated or even completed because of the greater cost efficiency, and the number of such studies is expected to increase, with several to be run in smaller labs^3-5^. The availability of fast, simple to use and accurate genotype calling pipelines is therefore of utmost importance.

GATK is the most frequently used pipeline^6^, but other pipelines have outperformed it in terms of the F_1_ statistic, precision, and recall. Specifically, DeepVariant^7^ won the first precisionFDA Truth Challenge for short-read sequencing in 2016 and had the highest accuracy of single nucleotide polymorphisms (SNPs). The winner of the second precisionFDA Truth Challenge for short-read sequencing in 2020 was Illumina’s DRAGEN^8^. It outperformed other pipelines^6,9-12^, in particular in difficult-to-call genomic regions^13^. It is, however, unclear which pipeline should be used for secondary analysis of WGS data, i.e., from fastq to vcf, in terms of both speed and accuracy. The best choice is challenging because recent pipeline comparisons relied on synthetic-diploid and simulated data^14^ or did not include the award-winning DeepVariant and/or DRAGEN pipelines^6,9,10,12,15^. Furthermore, several comparisons focused only on variant calling, the last step in the secondary analysis pipeline.

Since it is important to select the most appropriate secondary analysis pipeline for large-scale WGS efforts, we compared DRAGEN against GATK with BWA-MEM2 in the mapping & alignment steps (upstream pipeline), and DeepVariant and DRAGEN against GATK in the variant calling step (downstream pipeline, Figure 1). To assess the performance of the pipelines, we used samples from the genome in a bottle (GIAB) consortium as the reference truth set. We successfully sequenced one GIAB sample (NIST ID HG002) 70 times in different sequencing runs and one trio (NIST IDs HG002, HG003 and HG004) three times. These sequences allowed for the comparison of different pipelines against truth sets for single nucleotide variations (SNVs), insertions and deletions as well as complex variants (Indels) in simple-to-map, difficult-to-map, coding, and non-coding regions.

**Figure 1.**
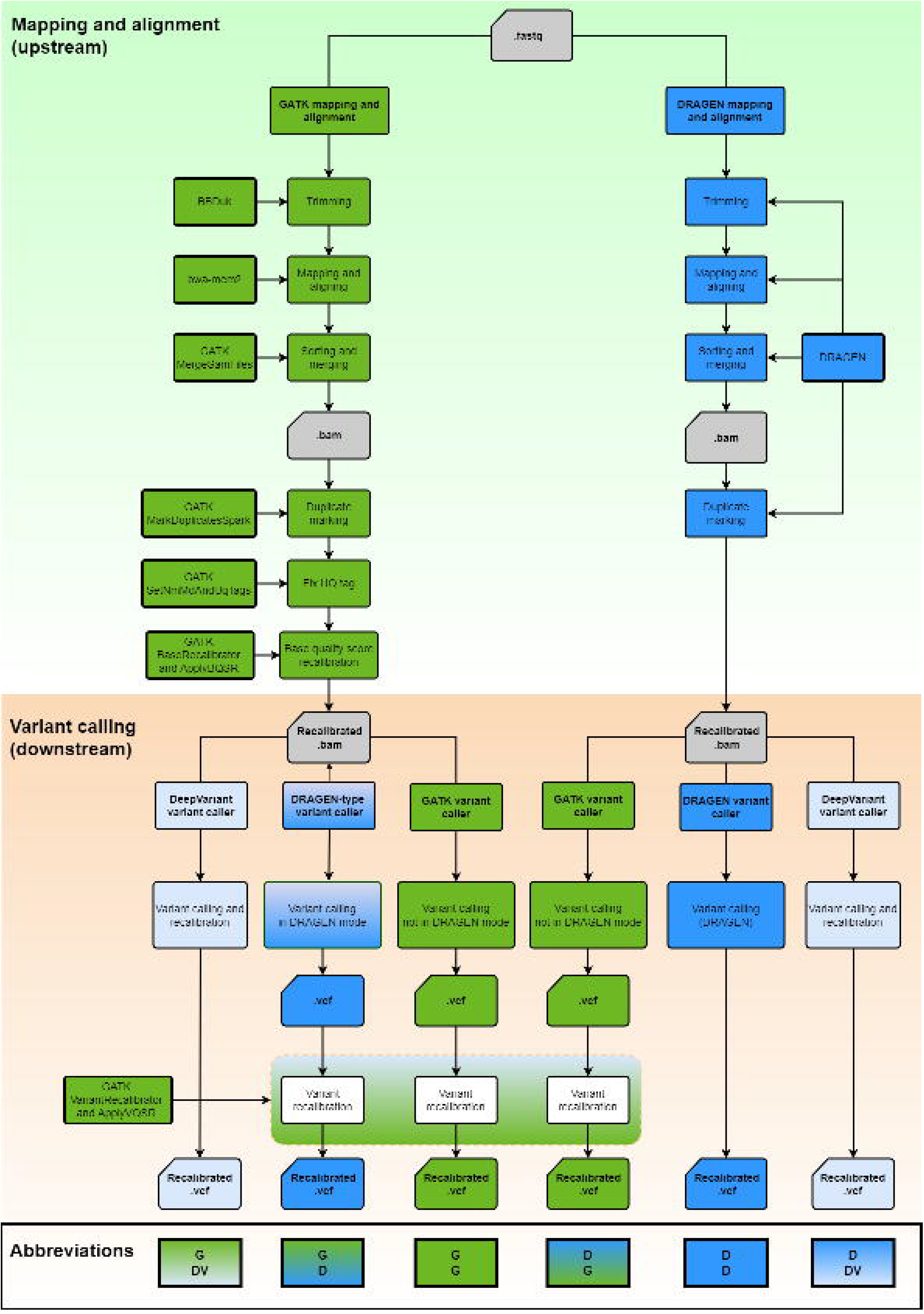
Flowchart of pipelines used in the benchmark analysis. The two upstream pipelines GATK and DRAGEN for mapping & alignment were used in conjunction with the four variant calling pipelines DRAGEN, DeepVariant, GATK Haplotypecaller in DRAGEN mode and GATK Haplotypecaller not in DRAGEN mode downstream.

## Results

### Run times

Total run times are displayed in Figure 2. The DRAGEN pipeline was fastest and required 36 ± 2 min (mean ± standard deviation) per sample; detailed comparisons are provided in Supplementary Table S1. The DRAGEN also showed the most homogeneous run time across all GIAB samples. The other pipelines were substantially slower with a minimum average run time ≥ 180 min. For example, DeepVariant variant calling following DRAGEN mapping & alignment required 256 ± 7 min. The long run time of DeepVariant was most likely caused by the single threading of the software in several of the computing steps. Run time for the upper part of the pipeline, i.e., up to the bam file, was 182 ± 36 min for GATK and 18 ± 1 min for the DRAGEN (Supplementary Figure S1). Run times for variant calling were 18 ± 1 min for the DRAGEN and 231 ± 16 min for DeepVariant (Supplementary Figure S1). The GATK Haplotypecaller in the DRAGEN mode took 189 ± 26 min to run, and 134 ± 20 min were required when GATK was run in the standard mode.

**Figure 2.**
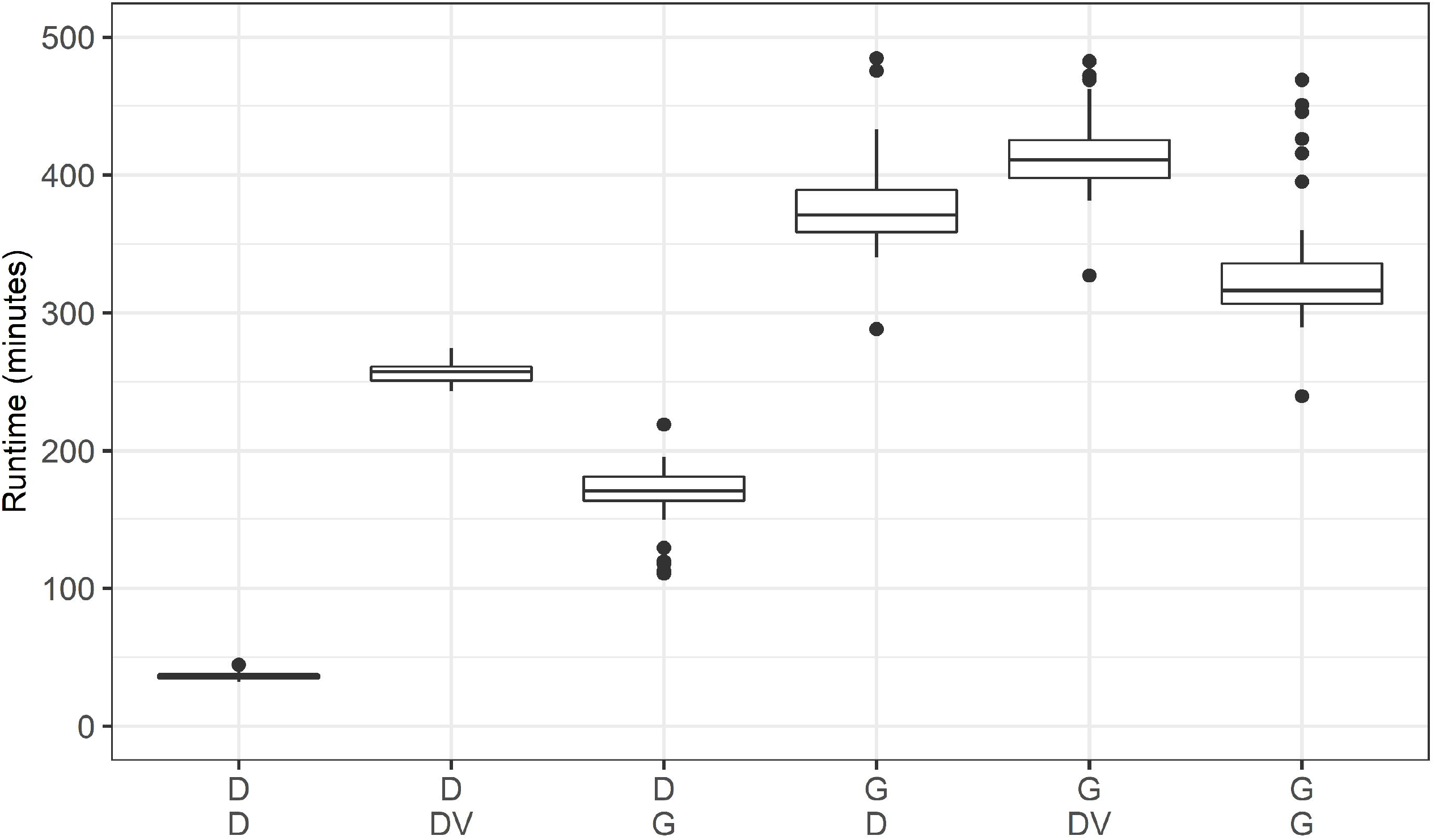
Runtime comparison of the 6 pipelines. The first row in the legend on the x-axis displays the approach used for mapping & alignment. The second row in the legend on the x-axis displays the variant caller used. In detail: D/D: DRAGEN for mapping & alignment and variant calling. D/DV: DRAGEN for mapping & alignment, DeepVariant for variant calling. D/G: DRAGEN for mapping & alignment, GATK with Haplotypecaller not in the DRAGEN mode for variant calling. G/D: GATK for mapping & alignment, GATK with Haplotypecaller in the DRAGEN mode for variant calling. G/DV: GATK for mapping & alignment, DeepVariant for variant calling. G/G: GATK for mapping & alignment, GATK without Haplotypecaller in the DRAGEN mode for variant calling.

### Called variants and Ti/Tv ratio

Table 1 shows the number of polymorphic sites that failed and passed the filtering step for HG002. Between 4,680,047 (GATK-DV) and 5,066,532 (DRAGEN-DRAGEN) variants passed the filtering step. Both DeepVariant variant calling pipelines had the highest number of positions with failed variants. When DRAGEN was used in the alignment step, on average >100,000 more filter passing variants were detected as compared to using GATK for mapping & alignment. Furthermore, when DRAGEN was used for variant calling, approximately 200,000 additional variants passed filtering as compared with DeepVariant or GATK in the variant calling (Table 1).

The transition-to-transversion (Ti/Tv) ratio is an important measure to assess the quality of SNV calling. After stringent quality control, WGS studies are expected to have a Ti/Tv ratio around 2.0 – 2.2^16^. For our GIAB samples, Ti/Tv ratios varied between 1.960 (DRAGEN-DRAGEN and DRAGEN-GATK) and 1.998 (DRAGEN-DeepVariant) prior to stringent quality control (Table 1).

### Benchmarking for SNVs and Indels

Figure 3 displays F_1_ scores, precision, and recall for SNVs (upper part A) and Indels (lower part B), respectively, for the 6 different pipelines after filtering based on chromosomes 20 to 22 of HG002. Both parts of the figure show substantially higher F_1_ scores when DRAGEN was used upstream compared to GATK. DeepVariant and DRAGEN performed similarly in the variant calling step when mapping & alignment was done with the DRAGEN. F_1_ scores and precision were slightly higher for SNVs with DeepVariant compared to DRAGEN, which has its basis in higher precision. In contrast, the recall for SNVs was lower for DeepVariant than for DRAGEN.

**Figure 3.**
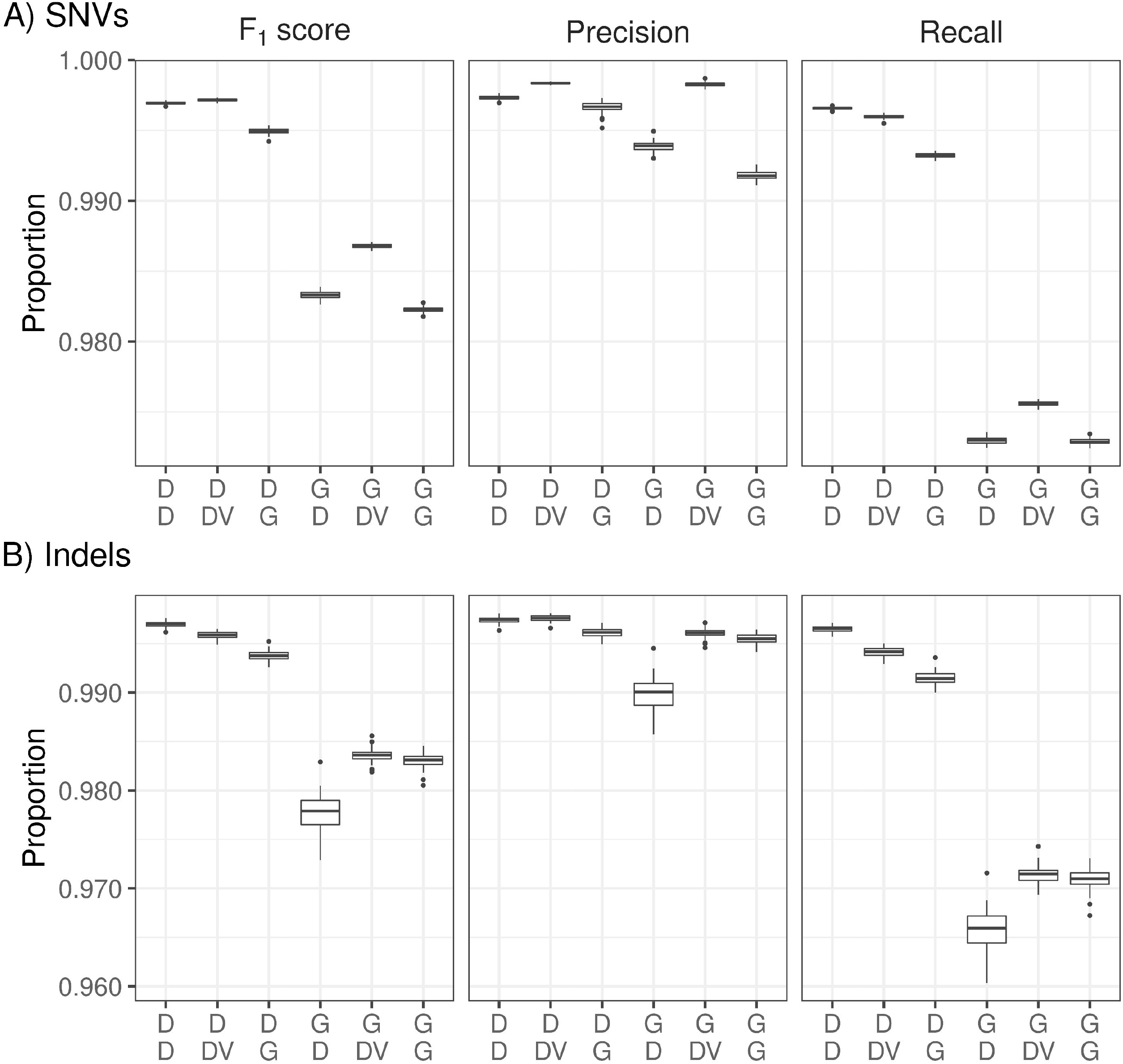
Percentages of F_1_ score, precision, and recall for single nucleotide variations (SNVs) (upper three panels) and Indels (lower three panels) for the 6 pipeline combinations, based on chromosomes 20 to 22 of the genome in a bottle sample HG003. Labels on the x-axis are defined in detail in Figure 2. D: DRAGEN; DV: DeepVariant; G: GATK.

In the case of Indels, the F_1_ score of the DRAGEN was higher than for DeepVariant, when the DRAGEN was used upstream. Supplementary Figure S2 confirms these findings using all autosomes for GIAB sample HG003. On this sample, the DRAGEN outperformed GATK with BWA-MEM2 upstream. The DRAGEN also slightly outperformed DeepVariant in the variant calling step on the F_1_ score for both SNVs and Indels. However, the precision of DeepVariant was still higher for DeepVariant compared to DRAGEN variant calling for both SNVs and Indels.

### Benchmarking for SNVs and Indels in simple-to-map (simple) and difficult-to-map (complex) regions

Results were similar for difficult-to-map (complex) and simple-to-map (simple) regions based on chromosomes 20 to 22 of HG002 (Figure 4). These regions have been defined for the truth set of the GIAB samples; for details, see Methods. For SNVs, F_1_ scores were lower in complex regions, when GATK was used upstream compared to any DRAGEN pipeline upstream. These differences in the F_1_ statistic were primarily caused by low recall values for GATK-based pipelines. However, even precision values were lower for SNVs in complex regions when GATK was used for mapping & alignment. In simple regions, precision was similar for all pipelines in case of SNVs, while recall was lower when GATK was used. Consequently, F_1_ scores were higher for all DRAGEN-based pipelines than for all GATK-based pipelines. A more detailed display for short, medium, and long insertions and deletions by regions is provided in Supplementary Figures S3 and S4, respectively.

**Figure 4.**
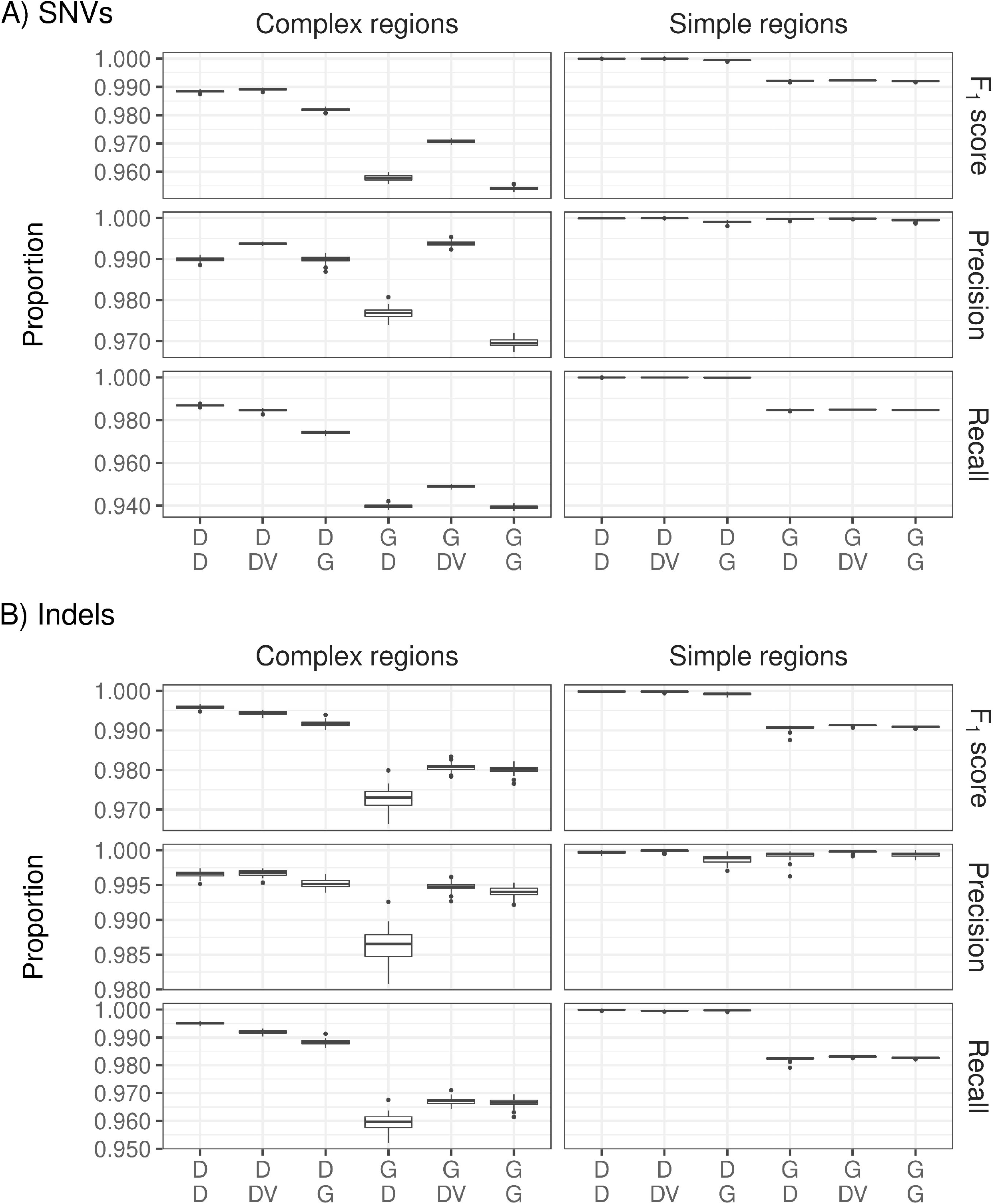
Percentages of F_1_ score, precision, and recall for complex-to-map (complex) and simple-to-map (simple) regions for single nucleotide variations (SNVs) (upper 6 panels) and Indels (lower 6 panels) for the 6 pipeline combinations, based on chromosomes 20 to 22 of the genome in a bottle sample HG003. Labels on the x-axis are defined in detail in Figure 2. D: DRAGEN; DV: DeepVariant; G: GATK.

### Benchmarking for SNVs and Indels in coding and non-coding regions

F_1_ scores were higher for all DRAGEN-based pipelines than for all GATK-based pipelines in both coding and non-coding regions (Figure *5*). DeepVariant and DRAGEN performed similarly, when DRAGEN was used upstream, with some minor advantages for the DRAGEN.

**Figure 5.**
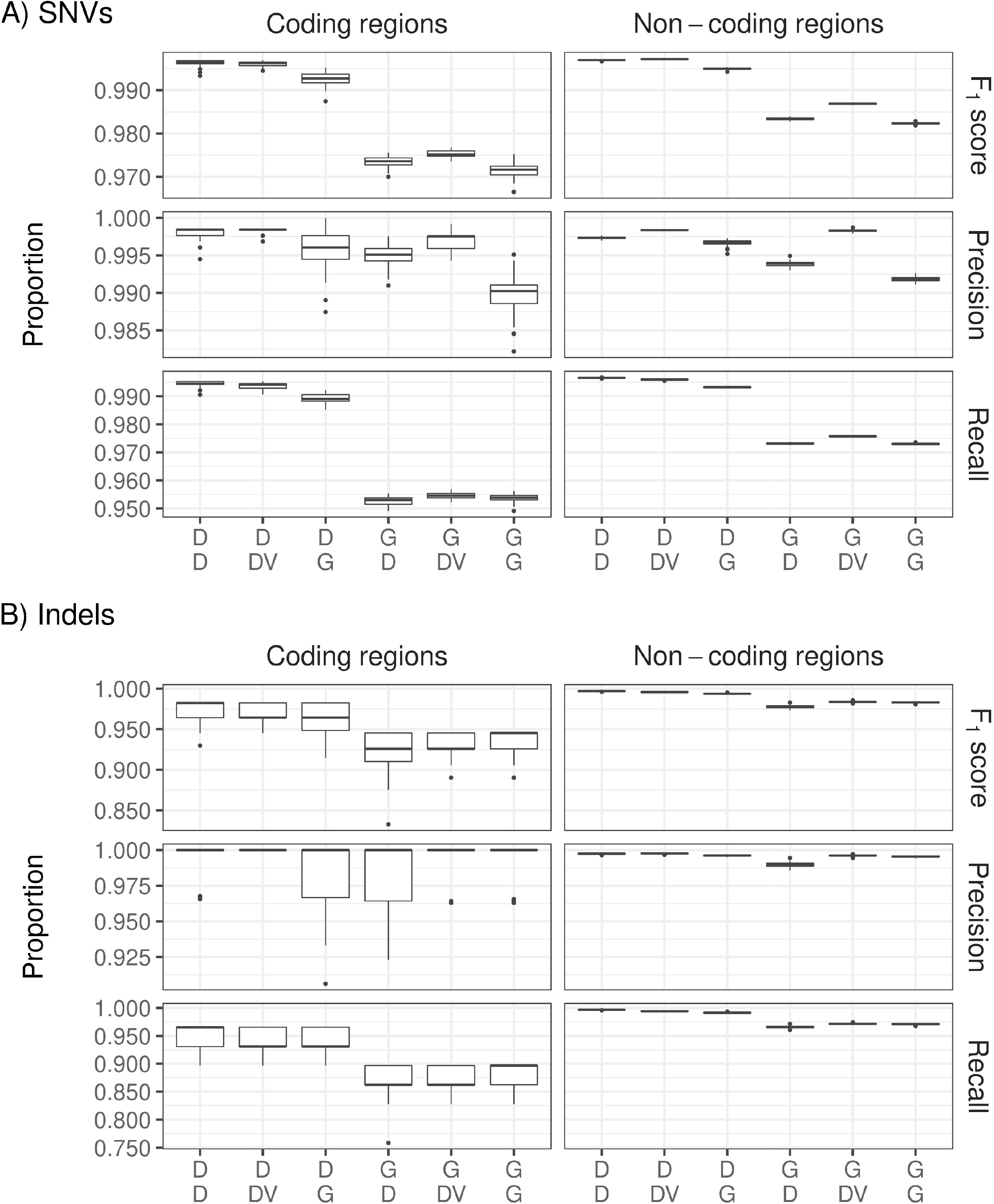
Percentages of F_1_ score, precision, and recall for coding and non-coding regions for single nucleotide variation (SNV) (upper 6 panels) and Indels (lower 6 panels) for the 6 pipeline combinations, based on chromosomes 20 to 22 of the genome in a bottle sample HG003. Labels on the x-axis were defined in detail in Figure 2. D: DRAGEN; DV: DeepVariant; G: GATK.

### Benchmarking using precision, recall and F_1_ scores for Indels of different size

Figure *6* shows F_1_ scores, precision, and recall for insertions (upper part) and deletions (lower part) of different sizes. Again, pipelines that used DRAGEN for mapping & alignment outperformed comparable GATK pipelines. Differences increased with the size of the insertions or deletions. In the variant calling step, DRAGEN overall showed a better performance than DeepVariant.

**Figure 6.**
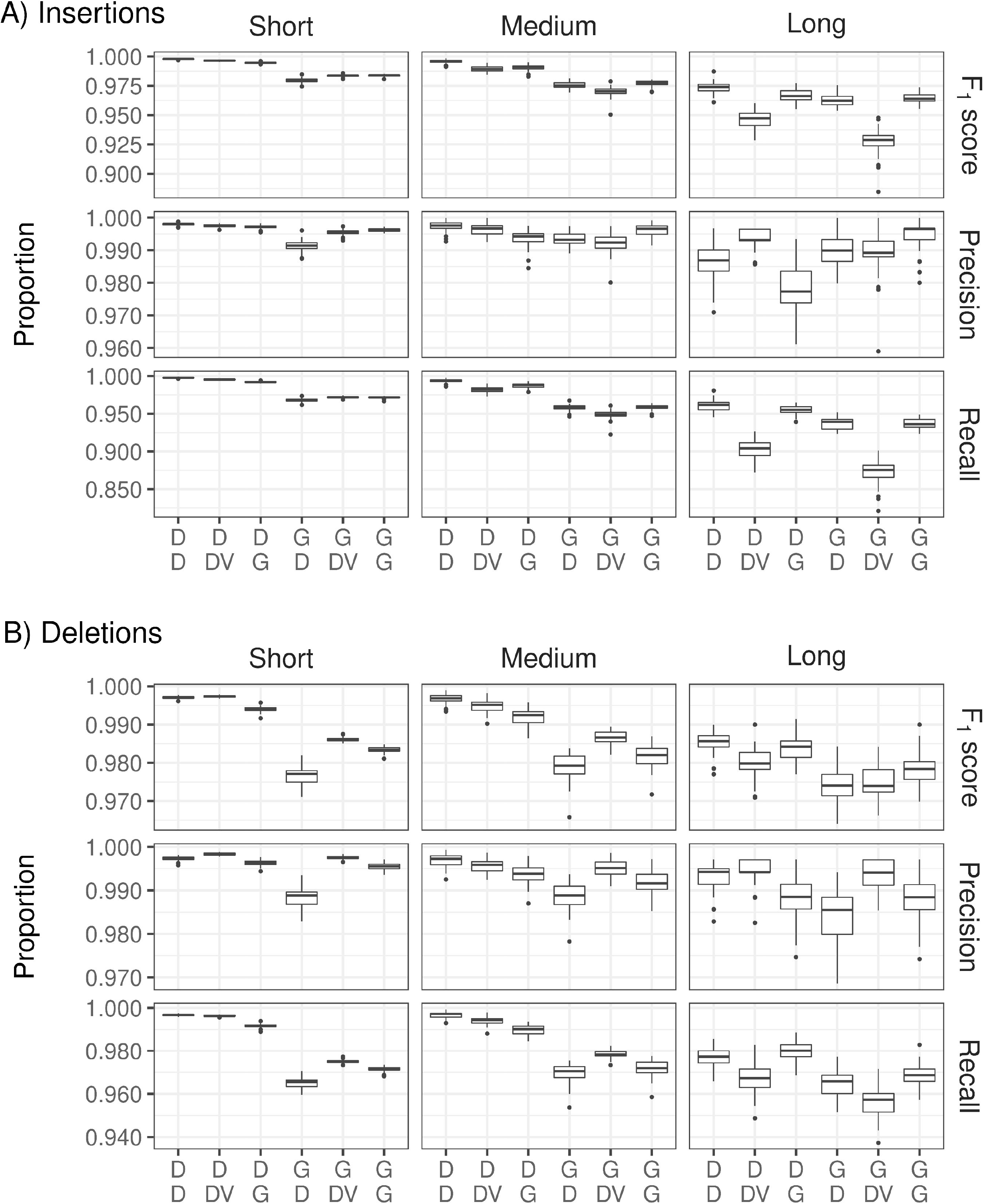
Percentages of F_1_ score, precision and recall for deletions (upper part A) and insertions (lower part B) for short (1-5 bp), medium (6-15 bp) and long (> 15 bp) insertions / deletions for the 6 pipeline combinations, based on chromosomes 20 to 22 of the genome in a bottle sample HG003. Labels on the x-axis are defined in detail in Figure 2. D: DRAGEN; DV: DeepVariant; G: GATK.

### Mendelian inheritance error fractions

The GIAB trio consisting in HG002, HG003, and HG004 was sequenced three times to estimate Mendelian inheritance error fractions. Figure 7 shows that Mendelian inheritance error fractions were lower for the in-built DRAGEN variant caller. Overall, the DRAGEN mapping & alignment outperformed GATK with BWA-MEM2 mapping & alignment.

**Figure 7:**
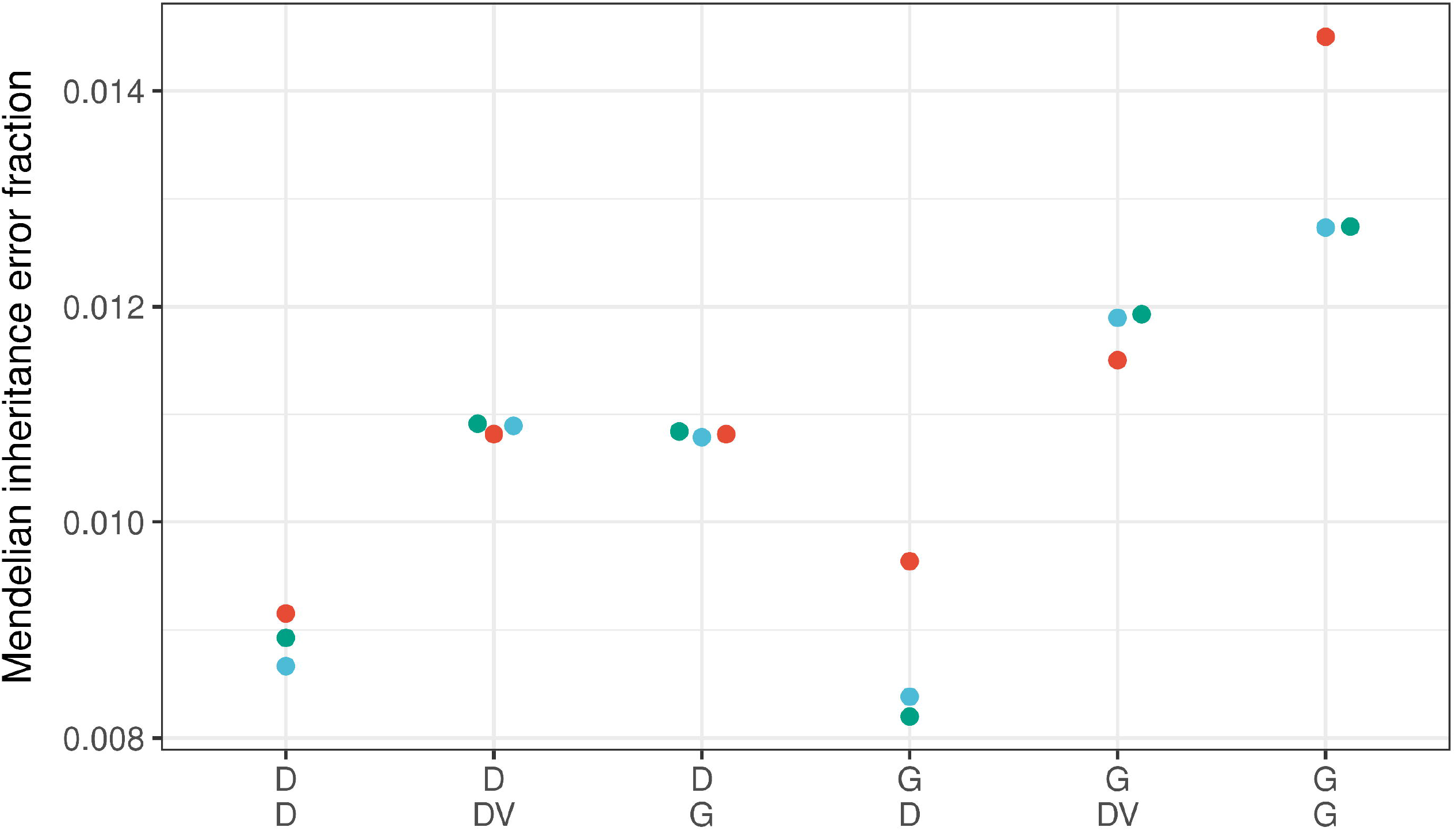
Mendelian inheritance error fractions. The GIAB trio consists of the genome in a bottle samples HG002, HG003, and HG004. They were sequenced three times, and Mendelian inheritance error fractions were estimated for chromosomes 20 to 22. Each dot represents the Mendelian inheritance error fraction for one trio. Labels on the x-axis are defined in detail in Figure 2.

## Discussion

Our empirical comparison of 6 different pipeline combinations for mapping & alignment and variant calling of WGS data demonstrated that mapping & alignment with DRAGEN had higher F_1_ scores, precision, and recall compared to GATK with BWA-MEM2. Specifically, the DRAGEN outperformed GATK in the upstream part of the pipeline for both SNVs and Indels, even when stratified by mapping complexity, coding or non-coding regions, and by Indel size. Differences were most pronounced for the recall, termed sensitivity in statistics, i.e., the probability to detect a true variant. In line with previous precisionFDA Truth Challenges and other work^8,9^, DeepVariant and DRAGEN performed well in the calling of both SNV and Indels. DeepVariant was optimized for SNV calling^7^, and it performed slightly better than the DRAGEN variant caller on SNVs. In contrast, the DRAGEN performed better than DeepVariant on Indels. We failed to confirm the findings of Hwang and colleagues^15^ who demonstrated an excellent performance of BWA-MEM for mapping/alignment when combined with the GATK Haplotypecaller.

Computational efficiency was another aspect of interest in our comparative study. The highest speed was observed for the DRAGEN. Our GIAB samples had an average coverage of approximately 35×, and the processing time per sample was 36 ± 2 min for both mapping/alignment and variant calling, which was substantially faster than GATK. Others considered “computational time the most important advantage of the DRAGEN platform”^14^. However, our pipeline comparison showed the high accuracy of the DRAGEN in the mapping & alignment step. We consider this to be even more important because it affects downstream association and functional annotation analyses. The aspect of accuracy is not only demonstrated by the repeated sequencing of the GIAB sample HG002 and the GIAB sample HG003 (Supplementary Figure S2) and their comparison with the truth set data, but also by the Mendelian inheritance errors detected in the repeatedly sequenced GIAB trio. On the one hand, DeepVariant had fewer Mendel errors than GATK-based variant calling on parent-offspring trios after GATK-based mapping & alginment, which is in line with the findings of Lin and colleagues^6^. On the other hand, when the DRAGEN was used for mapping & alignment, there was almost no difference between DeepVariant and GATK in the variant calling step. Both mapping & alignment approaches resulted in similar, specifically lowest Mendel errors when DRAGEN was used for variant calling. Results might differ when DeepTrio is used rather than DeepVariant for variant calling in trios.

Furthermore, variants that pass from variant callers are generally further filtered during gvcf joint calling steps. In the present work we have focused on the pipeline from fastq to single sample gvcf. Future work should investigate the effect of the different joint calling and gvcf filtering approaches.

One limitation of our trio analysis is that all subjects were treated individually. Inheritance-exploiting variant callers, such as DeepTrio^17^ were not taken into consideration. However, large WGS studies generally focus on the association analysis of unrelated subjects.

As different sequencers may have distinct error profiles^18^, one may consider the use of a single NovaSeq 6000 sequencer a limitation in the presented sequencing experiment, and it would be of interest to expand the sequencing to different platforms. However, the expensive platforms HiSeq X Ten and NovaSeq 6000 have low error rates and low levels of variation^18^. Furthermore, the NovaSeq 6000 sequencer is the only machine currently recommended by Illumina for large scale WGS. The just announced NovaSeq X series is not available on the market yet, and the HiSeq X Ten platform is not available anymore. Finally, the sequences were generated on a single NovaSeq 600 by only two lab technicians, the resulting sequencing data generated are as homogeneous as possible.

Another limitation might be that DeepVariant was trained by incorporating all GIAB samples but HG003 and all autosomal data but chromosomes 20 to 22^19^. For our evaluation, we restricted our analysis to chromosomes 20 to 22 of HG002 to prevent overfitting because these chromosomes were excluded from all training and model selection steps for all samples for DeepVariant. Sensitivity analyses involved GIAB sample HG003 (Supplementary Figure S2), for which all data were not included in the training of DeepVariant. Finally, we also analyzed all chromosomes of HG002 (data not shown). In all three cases, the performance of DeepVariant differed only marginally. Future studies might consider constructing a super learner in the variant calling step involving the two best-performing variant callers in this study, DeepVariant and DRAGEN.

## Conclusions

The DRAGEN mapper and aligner had higher accuracy than the GATK with BWA-MEM2 mapper and aligner. DeepVariant and DRAGEN performed similarly for SNV and Indel variant calling, and both outperformed GATK variant calling pipelines. The DRAGEN pipeline showed the lowest percentage of Mendelian inheritance errors and had the shortest execution times. Because of accuracy and speed, we recommend the use of the DRAGEN for secondary analysis of WGS data.

## Methods

### Sample preparation

We ordered the GIAB samples from the Coriell Institute (NA24385, NIST ID HG002; NA24149, NIST-ID HG003 and NA24143, NIST-ID HG004). DNA concentration was measured by Qubit.

The library was constructed according to Illumina TruSeq DNA PCR Free Library Prep protocol HT (Illumina Inc., San Diego, CA, USA) for whole genome sequencing. Briefly, the protocol steps were: (1) fragmentation of 1 μg genomic DNA to 350 bp inserts by Covaris LE220-plus, (2) cleanup of fragmented DNA, (3) repair ends, (4) removal of large and small DNA fragments, (5) 3’-end adenylation and (6) adapter ligation. The resulting library was quantified and quality-assessed with the iSeq100 (Illumina). The GIAB samples were sequenced with the NovaSeq 6000 platform (Illumina) using S4 flow cells with 300 cycles (2x 150 reads) and measured 2x to reach an average coverage of 35x.

### Variant calling pipelines

The raw sequencing files (base call file) were converted to fastq format and demultiplexed in a single step using Illumina’s bcl2fastq program on a single DRAGEN version 2^20^. First steps in secondary analysis, including mapping & alignment, sorting, duplicate marking, and base quality recalibration were done using the GATK pipeline (version 4.2.4.1)^21^ and the DRAGEN pipeline (version 3.8.4)^22^ (Figure 1). The GATK best practices workflow was applied^21^. Read trimming for the GATK pipeline was performed with BBDuk (version 38.90) using the following parameters: ktrim=r k=23 mink=11 hdist=1 tpe tbo threads=8 trimpolygright=10^23^. For the DRAGEN pipeline, the integrated soft read trimer with default parameters was used. The human reference genome hs38DH was used for mapping, which contains the primary assembly of GRCh38 plus ALT contigs, additional decoy contigs and HLA genes^24^. BWA-MEM2 2.2.1 was employed for mapping & alignment^25^, which is faster than BWA-MEM^25^. In the GATK base quality score recalibration step, the genome was split into 64 fractions for higher computational efficiency (one fraction per core). In the variant calling step, 25 fractions were used. Variant calling for SNVs and Indels was performed using the GATK HaplotypeCaller not in the DRAGEN mode, the GATK HaplotypeCaller in the DRAGEN mode^26^, DeepVariant v1.1.0^7^, and DRAGEN^22^. Alignment and variant calling algorithms were combined to 6 different pipelines (Figure 1): 1) GATK upstream plus DeepVariant downstream, 2) GATK upstream plus GATK HaplotypeCaller in DRAGEN mode downstream, 3) GATK upstream plus GATK HaplotypeCaller not in DRAGEN mode downstream, 4) DRAGEN upstream and downstream, 5) DRAGEN upstream plus DeepVariant downstream and 6) DRAGEN upstream plus GATK Haplotypecaller not in DRAGEN mode downstream. Variants with pure addition or removal are defined as insertions and deletions. For instance, A (REF) -> AT (ALT) is a one base insertion, ATTT (REF) -> AT (ALT) is a two base deletion. Variants with a length change between the REF and ALT allele which is not pure are defined as complex variants. ATT (REF) -> CTTT (ALT) would correspond to a complex variant with one base changed and one base inserted. Indels include insertions, deletions, and complex variants. To filter the variants produced by GATK Haplotypecaller, variant quality score recalibration (VQSR) was applied with the parameters recommended by GATK: For Indel recalibration, the Mills_and_1000G_gold_standard.indels.hg38 and Axiom_Exome_Plus.genotypes.all_populations.poly.hg38 datasets were used for training. For SNV recalibration, the Hapmap_3.3.hg38, 1000G_omni2.5.hg38 and the 1000G_phase1.snps.high_confidence.hg38 datasets were used for training. The dbSNP_v154_GRch38.p12 was used as truth set in both cases. For DeepVariant, default filtering parameters were used. For DRAGEN, default filtering parameters were used, except the QUAL threshold was changed from < 10.41 for SNVs and < 7.83 for Indels to <5.0 for both SNVs and Indels upon recommendation from the Illumina bioinformatics team (personal communication).vcf files were used for analysis. For each trio, the gvcf files created from each pipeline were jointly called with GLNexus (version 1.4.1)^27^, except for the DRAGEN mapping/alignment and variant calling pipeline. Specifically, DRAGEN treats non-PAR regions of chromosome X as haploid, while GATK and DeepVariant treat them as diploid. So far, DRAGEN gvcf files are not supported by GLNexus. DRAGEN gvcf files were therefore jointly called with GenomicsDBImport and GenotypeVcfs, both part of GATK. The Mendelian inheritance error was calculated with bcftools mendelian plugin (version 1.11)^28^.

### Computing environment and resources

Analyses were run on a local high-performance computing cluster at Cardio-CARE (Davos, CH). Each compute node is equipped with 2 AMD EPYC 7742 CPUs, 2 TB RAM, approximately 11 TB NVMe, and CentOS 8 operating system. For pipeline comparisons, each pipeline was given 64 cores as in Ref. ^14^. Two DRAGEN Servers v3 were used for the DRAGEN pipeline.

### Statistics

Precision, recall and F_1_ scores were estimated for SNVs and Indels using all regions and regions stratified by difficult-to-map, simple-to-map and coding and non-coding regions. All comparisons were made against the GIAB truth set^29,30^, available from ftp.ncbi.nlm.nih.gov/giab/ftp/release/AshkenazimTrio/. Mappability of Indels was stratified by Indel size (1-5 bp, 6-15 bp, > 15 bp). Ti/Tv ratios were calculated to indicate potential sequencing error. The target sensitivity was varied and fixed to 99.75% for Indels and 99.95% for SNVs. Figures for varying target sensitivities are shown in Supplementary Figure S6. Results from the 70 GIAB sample sequencing repetitions are summarized in boxplots. Performance metrics were calculated using hap.py v0.3.14 with vcfeval engine, available from github.com/Illumina/hap.py. The numbers of passed/failed variants, SNVs, insertions and deletions for each pipeline were calculated using RTG tools (version 3.12.1)^31^. The statistical analysis was performed with R (version 4.1.1) to calculate mean differences and 95% confidence intervals between runtimes (Supplementary Table S1) and performance metrics (Supplementary Table S2).

### Genome regions for stratification analyses

To compare the performance of the calling algorithms in simple-to-map, sometimes termed easy-to-map, and difficult-to-map regions of the human genome, we stratified the data in genome-specific regions by using complexandSVs_alldifficultregions and notin_complexandSVs_alldifficultregions^30^. Stratification for coding regions was based on notinrefseq_cds and refseq_cds stratifications. These annotations are available at github.com/genome-in-a-bottle/genome-stratifications.

## Data availability

The datasets generated during and/or analyzed during the current study are available from the corresponding author on reasonable request for collaborative projects.

## Acknowledgements

We gratefully acknowledge funding of the whole genome sequencing study by the Kühne Foundation. Raphael Twerenbold holds a professorship funded by the Kühne Foundation. Tanja Zeller is supported by the German Center for Cardiovascular Research (DZHK e.V.) (81×2710170, partner site project).

## Author contributions

A.T., A.Z. and R.B. designed this study. A.T. and R.B. performed the bioinformatic and statistical analysis. D.A.G. performed the first quality analysis of the genome data. M.Z. and H.M. supervised the laboratory work. S.B. and T.Z. developed the concept of the whole genome sequencing study. A.Z. and R.B. wrote the paper. All authors commented on the manuscript.

## Additional Information

Supplementary Information. The online version contains supplementary material available at https://doi.org/XXX

## Competing Interests

A.T., A.Z., R.B. and S.B. are employees of Cardio-CARE, a 100% daughter of the Kühne Foundation.

## Table headings

Table 1. Comparison of variant counts for the 6 pipelines with HG002 sequenced 70 times in different runs. Displayed are means and standard deviations (in parenthesis).

## Notes

### Competing Interest Statement

A.T., A.Z., R.B. and S.B. are employees of Cardio-CARE, a 100% daughter of the Kuehne Foundation.

### Summary of Updates

All parts of the manuscript have undergone a revision.

