## Supplementary Material 1 for "Comparison of calling pipelines for whole genome sequencing: an empirical study demonstrating the importance of mapping and alignment"

**Supplementary Figures**

Figure S1: Runtime comparisons. Left: between the DRAGEN and GATK pipeline for mapping & alignment. Right: between DRAGEN, DeepVariant (DV), GATK mapping & alignment plus GATK Haplotypecaller in the DRAGEN mode (GATK / DRAGEN), and GATK mapping & alignment plus GATK Haplotypecaller not in the DRAGEN mode (GATK / GATK).

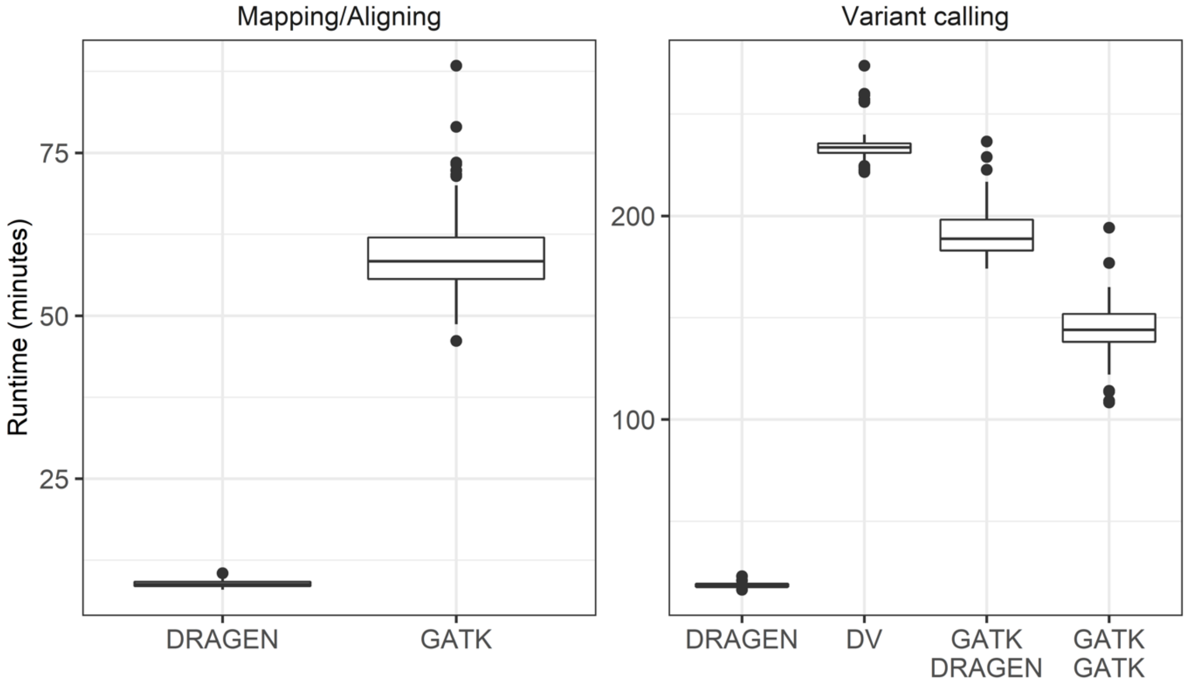

Figure S2: Performance evaluation of all 6 pipelines combination for single nucleotide variations (SNVs) and Indels, based on all autosomes of the genome in a bottle sample HG003. D: DRAGEN; DV: DeepVariant; G: GATK. First row in legend on x-axis displays approach used for mapping & alignment. Second row in legend on x-axis displays variant caller used. In detail: D/D: DRAGEN for mapping & alignment and variant calling. D/DV: DRAGEN for mapping & alignment, DeepVariant for variant calling. D/G: DRAGEN for mapping & alignment, GATK with Haplotypecaller not in the DRAGEN mode for variant calling. G/D: GATK for mapping & alignment, GATK with Haplotypecaller in the DRAGEN mode for variant calling. G/DV: GATK for mapping & alignment, DeepVariant for variant calling. G/G: GATK for mapping & alignment, GATK without Haplotypecaller in the DRAGEN mode for variant calling.

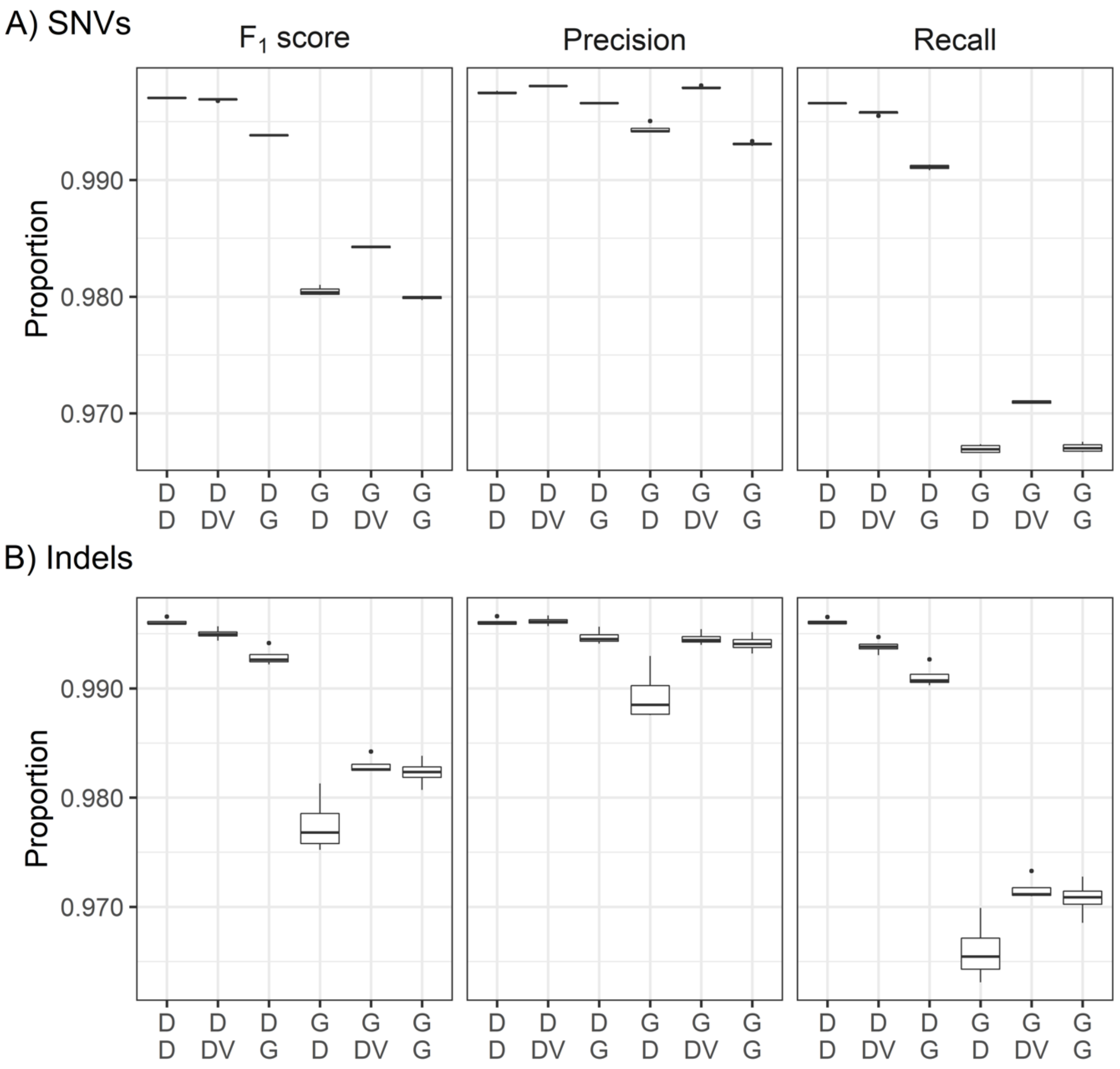

Figure S3: Percentages of $F_1$ score, precision, and recall for insertions in complex regions (upper part A) and insertions in simple regions (lower part B) for short (1-5 bp), medium (6-15 bp), and long (> 15 bp) insertions for the 6 pipeline combinations, based on chromosomes 20 to 22 of the genome in a bottle sample HG002. Labels on the x-axis are defined in detail in Supplementary Figure S3.

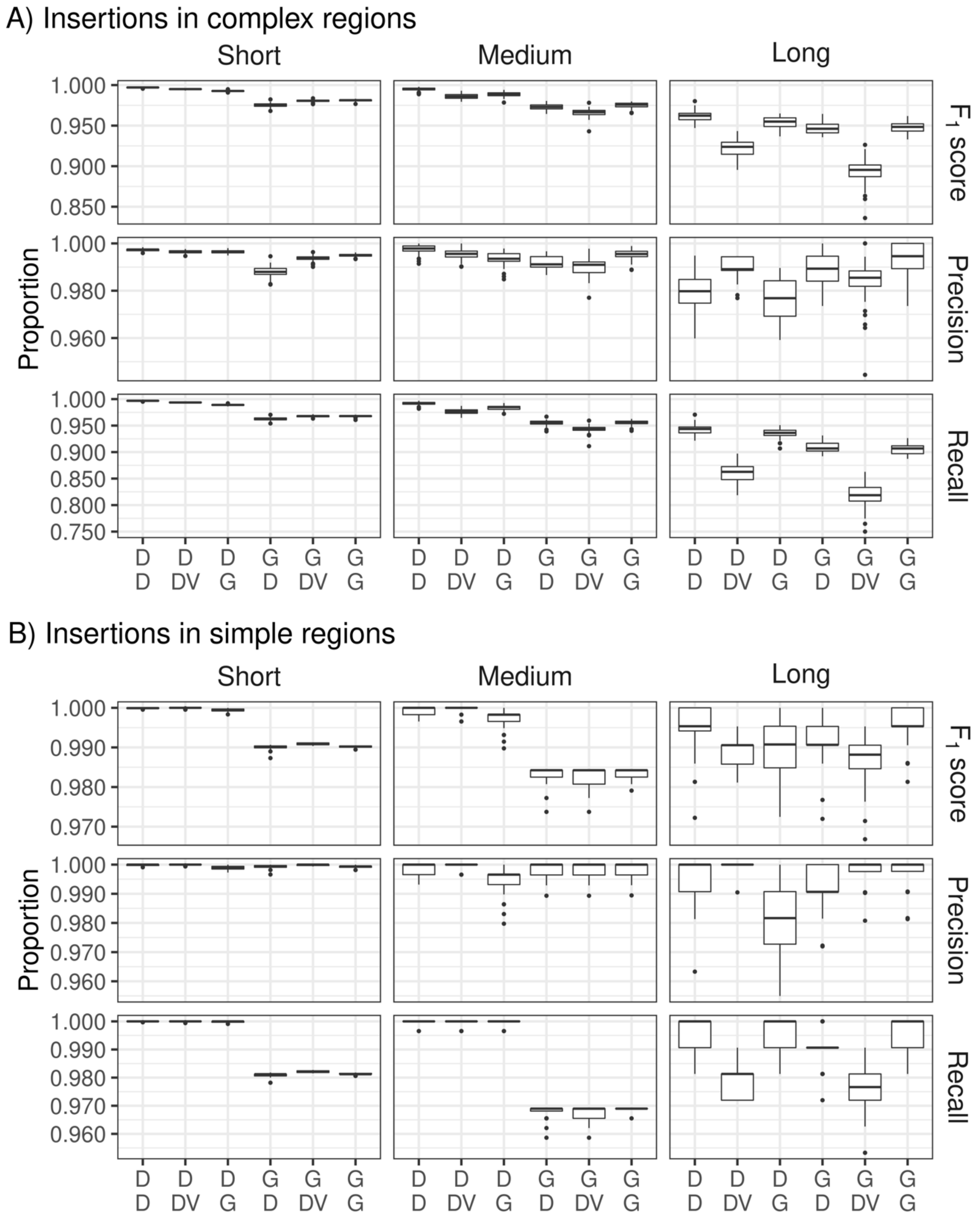

Figure S4: Percentages of $F_1$ score, precision, and recall for deletions in complex regions (upper part A) and deletions in simple regions (lower part B) for short (1-5 bp), medium (6-15 bp), and long (> 15 bp) deletions for the 6 pipeline combinations, based on chromosomes 20 to 22 of the genome in a bottle sample HG002. Labels on the x-axis are defined in detail in Supplementary Figure S3.

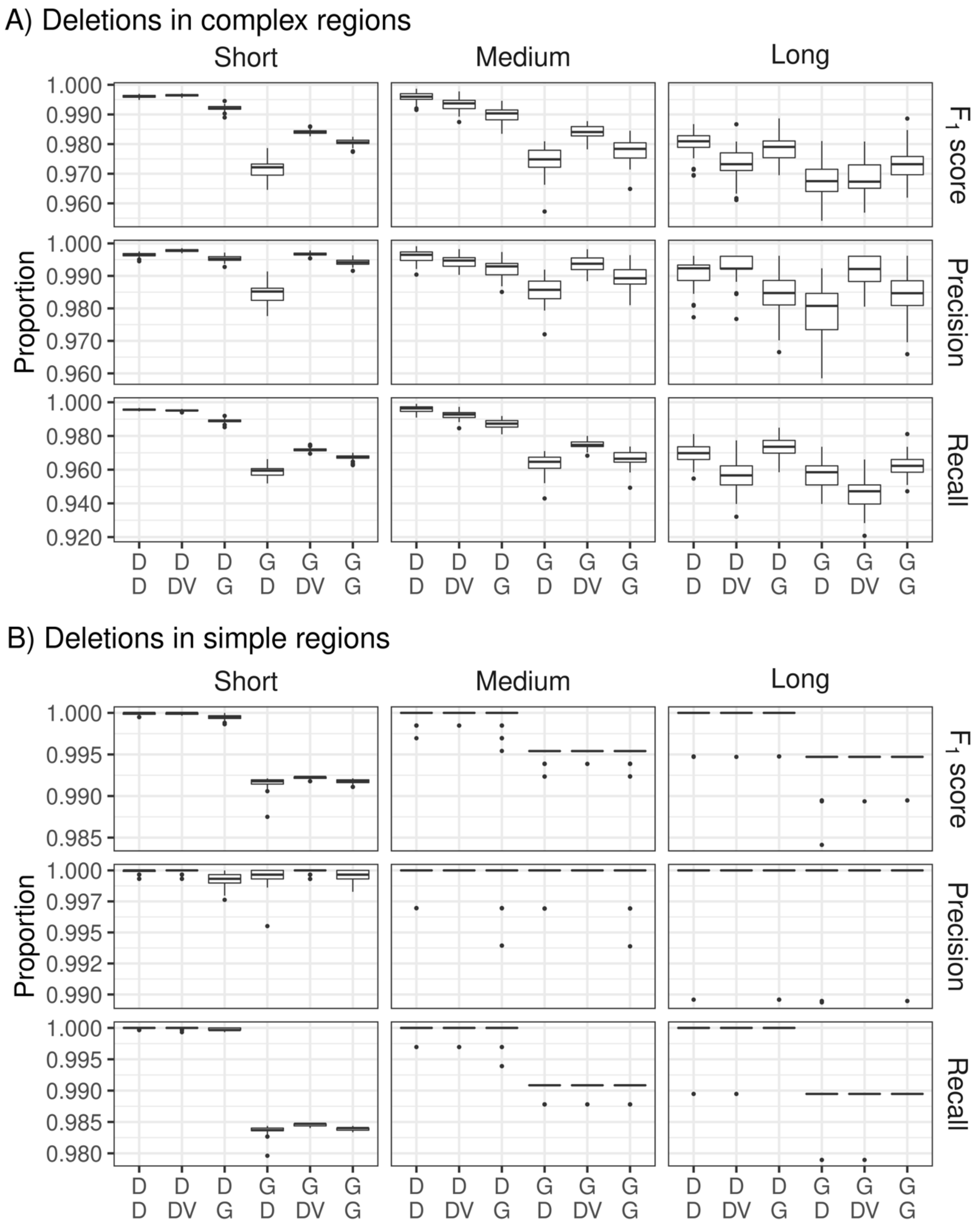

Figure S5: Performance evaluation of three pipelines for varying target sensitivities using the genome in a bottle sample HG002. Red: GATK Haplotypecaller (in DRAGEN mode, based on the bam file obtained from GATK with BWA-MEM2 mapping & alignment). Blue: GATK Haplotypecaller (not in DRAGEN mode, based on the bam file obtained from DRAGEN). Green: GATK Haplotypecaller (not in DRAGEN mode, based on the bam file obtained from GATK with BWA-MEM2 mapping & alignment). SNV: single nucleotide variation.

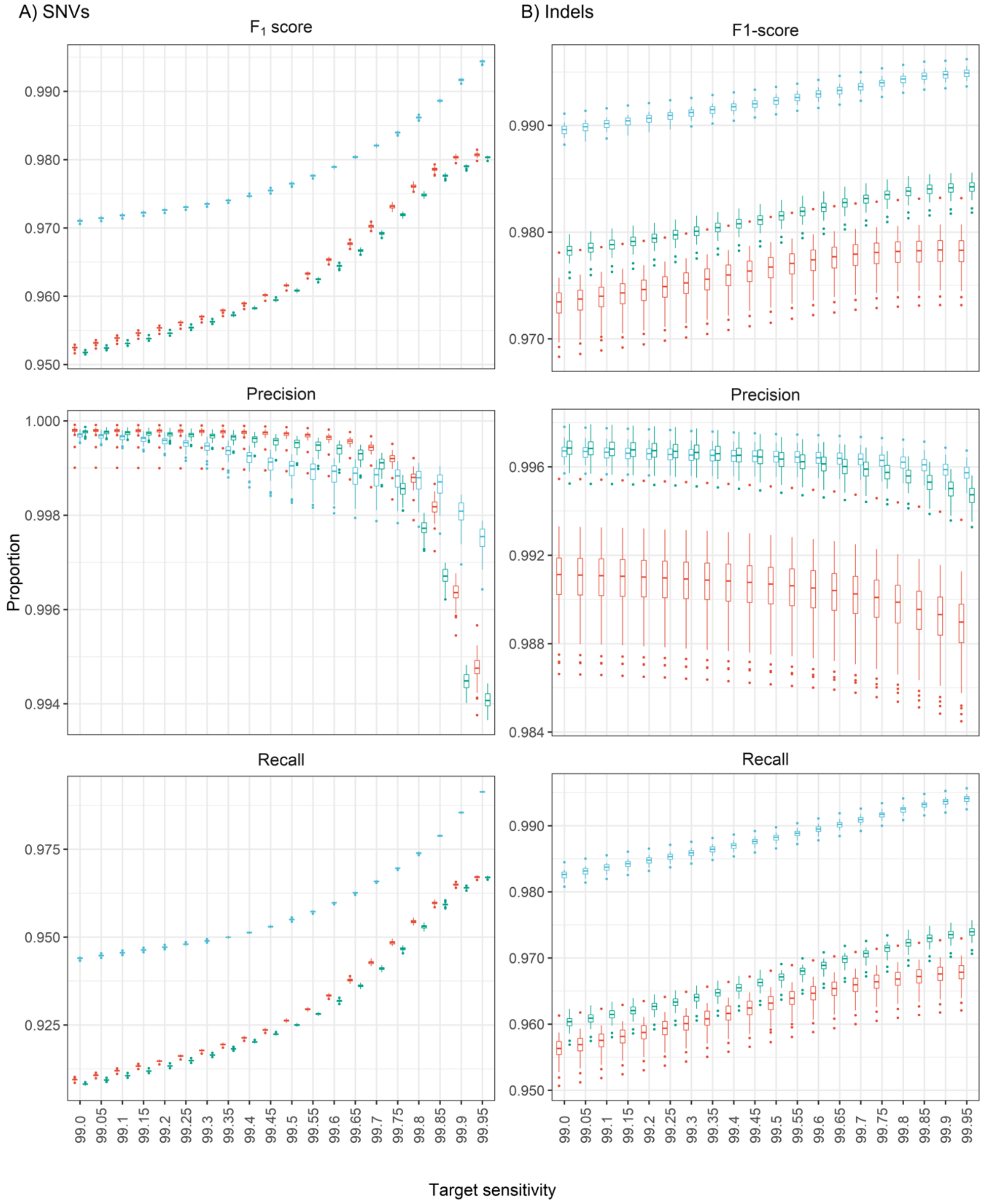

Table S1: Summary table of mean differences and 95% confidence intervals between the runtime of all 15 pipeline combinations.

| Pipeline 1 | Pipeline 2 | Mean difference | CI low | CI high |
|---|---|---|---|---|
| D-D | D-DV | -220 | -222 | -218 |
| D-D | D-G | -134 | -138 | -129 |
| D-D | G-D | -347 | -356 | -337 |
| D-D | G-DV | -385 | -394 | -376 |
| D-D | G-G | -291 | -300 | -281 |
| D-DV | D-G | 86 | 82 | 90 |
| D-DV | G-D | -127 | -136 | -117 |
| D-DV | G-DV | -165 | -174 | -156 |
| D-DV | G-G | -70 | -80 | -61 |
| D-G | G-D | -213 | -222 | -203 |
| D-G | G-DV | -251 | -261 | -242 |
| D-G | G-G | -157 | -166 | -148 |
| G-D | G-DV | -38 | -43 | -34 |
| G-D | G-G | 56 | 54 | 58 |
| G-DV | G-G | 95 | 90 | 100 |

Table S2: Summary table of mean differences and 95% confidence intervals of the performance for $F_1$ score, recall and precision for SNVs and Indels in simple, complex, coding, and non-coding and all regions for chromosomes 20 to 22 of HG002.

| Type | Region | Metric | Pipeline 1 | Pipeline 2 | Mean difference | CI low | CI high |
|---|---|---|---|---|---|---|---|
| Indel | All regions | Precision | D-D | D-DV | -0.000690655882 | -0.0009643992908 | -0.000416912474 |
| Indel | All regions | Precision | D-D | D-G | 0.002252808824 | 0.0019625243723 | 0.002543093275 |
| Indel | All regions | Precision | D-D | G-D | 0.004013229412 | 0.0035957459520 | 0.004430712872 |
| Indel | All regions | Precision | D-D | G-DV | 0.000710238235 | 0.0004093939092 | 0.001011082561 |
| Indel | All regions | Precision | D-D | G-G | 0.000862525000 | 0.0005085112640 | 0.001216538736 |
| Indel | All regions | Precision | D-DV | D-G | 0.002943464706 | 0.0025266007336 | 0.003360328678 |
| Indel | All regions | Precision | D-DV | G-D | 0.004703885294 | 0.0043473942019 | 0.005060376386 |
| Indel | All regions | Precision | D-DV | G-DV | 0.001400894118 | 0.0011946264648 | 0.001607161771 |
| Indel | All regions | Precision | D-DV | G-G | 0.001553180882 | 0.0013072836074 | 0.001799078157 |
| Indel | All regions | Precision | D-G | G-D | 0.001760420588 | 0.0012652287796 | 0.002255612397 |
| Indel | All regions | Precision | D-G | G-DV | -0.001542570588 | -0.0019467069815 | -0.001138434195 |
| Indel | All regions | Precision | D-G | G-G | -0.001390283824 | -0.0018368468431 | -0.000943720804 |
| Indel | All regions | Precision | G-DV | G-G | 0.000152286765 | -0.0001830383438 | 0.000487611873 |

| Type | Region | Metric | Pipeline 1 | Pipeline 2 | Mean difference | CI low | CI high |
|---|---|---|---|---|---|---|---|
| Indel | All regions | Recall | D-D | D-DV | 0.008774636765 | 0.0074499026287 | 0.010099370901 |
| Indel | All regions | Recall | D-D | D-G | 0.003233710294 | 0.0029278131103 | 0.003539607478 |
| Indel | All regions | Recall | D-D | G-D | 0.019208885294 | 0.0181387143172 | 0.020279056271 |
| Indel | All regions | Recall | D-D | G-DV | 0.024468611765 | 0.0225395681580 | 0.026397655371 |
| Indel | All regions | Recall | D-D | G-G | 0.016968577941 | 0.0159821861907 | 0.017954969692 |
| Indel | All regions | Recall | D-DV | D-G | -0.005540926471 | -0.0068302393304 | -0.004251613611 |
| Indel | All regions | Recall | D-DV | G-D | 0.010434248529 | 0.0089207267673 | 0.011947770292 |
| Indel | All regions | Recall | D-DV | G-DV | 0.015693975000 | 0.0147614351782 | 0.016626514822 |
| Indel | All regions | Recall | D-DV | G-G | 0.008193941176 | 0.0067889611576 | 0.009598921195 |
| Indel | All regions | Recall | D-G | G-D | 0.015975175000 | 0.0151010919445 | 0.016849258055 |
| Indel | All regions | Recall | D-G | G-DV | 0.021234901471 | 0.0194227590430 | 0.023047043898 |
| Indel | All regions | Recall | D-G | G-G | 0.013734867647 | 0.0129620076198 | 0.014507727674 |
| Indel | All regions | Recall | G-D | G-DV | 0.005259726471 | 0.0036747244094 | 0.006844728532 |
| Indel | All regions | Recall | G-D | G-G | -0.002240307353 | -0.0024887910494 | -0.001991823656 |

| Type | Region | Metric | Pipeline 1 | Pipeline 2 | Mean difference | CI low | CI high |
|---|---|---|---|---|---|---|---|
| Indel | All regions | Recall | G-DV | G-G | -0.007500033824 | -0.0090134926434 | -0.005986575004 |
| Indel | All regions | F1-score | D-D | D-DV | 0.006094848739 | 0.0052388150430 | 0.006950882436 |
| Indel | All regions | F1-score | D-D | D-G | 0.003914892857 | 0.0036356668309 | 0.004194118883 |
| Indel | All regions | F1-score | D-D | G-D | 0.016795546218 | 0.0163320229564 | 0.017259069481 |
| Indel | All regions | F1-score | D-D | G-DV | 0.018666602941 | 0.0175074765185 | 0.019825729364 |
| Indel | All regions | F1-score | D-D | G-G | 0.012972798319 | 0.0125471613147 | 0.013398435324 |
| Indel | All regions | F1-score | D-DV | D-G | -0.002179955882 | -0.0029794073372 | -0.001380504427 |
| Indel | All regions | F1-score | D-DV | G-D | 0.010700697479 | 0.0095614374196 | 0.011839957538 |
| Indel | All regions | F1-score | D-DV | G-DV | 0.012571754202 | 0.0120680707401 | 0.013075437663 |
| Indel | All regions | F1-score | D-DV | G-G | 0.006877949580 | 0.0058279764663 | 0.007927922693 |
| Indel | All regions | F1-score | D-G | G-D | 0.012880653361 | 0.0123813302231 | 0.013379976500 |
| Indel | All regions | F1-score | D-G | G-DV | 0.014751710084 | 0.0136688064546 | 0.015834613713 |
| Indel | All regions | F1-score | D-G | G-G | 0.009057905462 | 0.0086500447700 | 0.009465766154 |
| Indel | All regions | F1-score | G-D | G-DV | 0.001871056723 | 0.0005037095782 | 0.003238403867 |

| Type | Region | Metric | Pipeline 1 | Pipeline 2 | Mean difference | CI low | CI high |
|---|---|---|---|---|---|---|---|
| Indel | All regions | F1-score | G-D | G-G | -0.003822747899 | -0.0041104780635 | -0.003535017735 |
| Indel | All regions | F1-score | G-DV | G-G | -0.005693804622 | -0.0069584617027 | -0.004429147541 |
| Indel | Complex regions | Precision | D-D | D-DV | -0.000728613235 | -0.0010802008097 | -0.000377025661 |
| Indel | Complex regions | Precision | D-D | D-G | 0.002305104412 | 0.0019707834268 | 0.002639425397 |
| Indel | Complex regions | Precision | D-D | G-D | 0.005108085294 | 0.0045302464963 | 0.005685924092 |
| Indel | Complex regions | Precision | D-D | G-DV | 0.001090097059 | 0.0006808887271 | 0.001499305391 |
| Indel | Complex regions | Precision | D-D | G-G | 0.001006380882 | 0.0005259458853 | 0.001486815879 |
| Indel | Complex regions | Precision | D-DV | D-G | 0.003033717647 | 0.0025698561025 | 0.003497579192 |
| Indel | Complex regions | Precision | D-DV | G-D | 0.005836698529 | 0.0053516764273 | 0.006321720631 |
| Indel | Complex regions | Precision | D-DV | G-DV | 0.001818710294 | 0.0015344893771 | 0.002102931211 |
| Indel | Complex regions | Precision | D-DV | G-G | 0.001734994118 | 0.0013902220990 | 0.002079766136 |
| Indel | Complex regions | Precision | D-G | G-D | 0.002802980882 | 0.0021833698896 | 0.003422591875 |
| Indel | Complex regions | Precision | D-G | G-DV | -0.001215007353 | -0.0016901510576 | -0.000739863648 |
| Indel | Complex regions | Precision | D-G | G-G | -0.001298723529 | -0.0018191311449 | -0.000778315914 |

| Type | Region | Metric | Pipeline 1 | Pipeline 2 | Mean difference | CI low | CI high |
|---|---|---|---|---|---|---|---|
| Indel | Complex regions | Precision | G-D | G-DV | -0.004017988235 | -0.0045922955377 | -0.003443680933 |
| Indel | Complex regions | Precision | G-D | G-G | -0.004101704412 | -0.0044658204265 | -0.003737588397 |
| Indel | Complex regions | Precision | G-DV | G-G | -0.000083716176 | -0.0005561120034 | 0.000388679650 |
| Indel | Complex regions | Recall | D-D | D-DV | 0.012006510294 | 0.0101589686952 | 0.013854051893 |
| Indel | Complex regions | Recall | D-D | D-G | 0.004434704412 | 0.0040161819637 | 0.004853226860 |
| Indel | Complex regions | Recall | D-D | G-D | 0.022291333824 | 0.0210387095875 | 0.023543958060 |
| Indel | Complex regions | Recall | D-D | G-DV | 0.029913170588 | 0.0272324364741 | 0.032593904702 |
| Indel | Complex regions | Recall | D-D | G-G | 0.019618007353 | 0.0184446984565 | 0.020791316249 |
| Indel | Complex regions | Recall | D-DV | D-G | -0.007571805882 | -0.0093475772594 | -0.005796034505 |
| Indel | Complex regions | Recall | D-DV | G-D | 0.010284823529 | 0.0083957306257 | 0.012173916433 |
| Indel | Complex regions | Recall | D-DV | G-DV | 0.017906660294 | 0.0167620193352 | 0.019051301253 |
| Indel | Complex regions | Recall | D-DV | G-G | 0.007611497059 | 0.0059121434600 | 0.009310850658 |
| Indel | Complex regions | Recall | D-G | G-D | 0.017856629412 | 0.0168734537920 | 0.018839805031 |
| Indel | Complex regions | Recall | D-G | G-DV | 0.025478466176 | 0.0229575945531 | 0.027999337800 |

| Type | Region | Metric | Pipeline 1 | Pipeline 2 | Mean difference | CI low | CI high |
|---|---|---|---|---|---|---|---|
| Indel | Complex regions | Recall | D-G | G-G | 0.015183302941 | 0.0143072226837 | 0.016059383199 |
| Indel | Complex regions | Recall | G-D | G-DV | 0.007621836765 | 0.0053884645092 | 0.009855209020 |
| Indel | Complex regions | Recall | G-D | G-G | -0.002673326471 | -0.0030288242688 | -0.002317828672 |
| Indel | Complex regions | Recall | G-DV | G-G | -0.010295163235 | -0.0123735682222 | -0.008216758248 |
| Indel | Complex regions | F1-score | D-D | D-DV | 0.008698913866 | 0.0074371439759 | 0.009960683755 |
| Indel | Complex regions | F1-score | D-D | D-G | 0.004824123950 | 0.0044795472005 | 0.005168700699 |
| Indel | Complex regions | F1-score | D-D | G-D | 0.019904638655 | 0.0193484059792 | 0.020460871332 |
| Indel | Complex regions | F1-score | D-D | G-DV | 0.023428159664 | 0.0216270708862 | 0.025229248442 |
| Indel | Complex regions | F1-score | D-D | G-G | 0.015127084034 | 0.0146379329445 | 0.015616235123 |
| Indel | Complex regions | F1-score | D-DV | D-G | -0.003874789916 | -0.0050873340159 | -0.002662245816 |
| Indel | Complex regions | F1-score | D-DV | G-D | 0.011205724790 | 0.0096627084626 | 0.012748741117 |
| Indel | Complex regions | F1-score | D-DV | G-DV | 0.014729245798 | 0.0139917204866 | 0.015466771110 |
| Indel | Complex regions | F1-score | D-DV | G-G | 0.006428170168 | 0.0050718808783 | 0.007784459458 |
| Indel | Complex regions | F1-score | D-G | G-D | 0.015080514706 | 0.0145112224965 | 0.015649806915 |

| Type | Region | Metric | Pipeline 1 | Pipeline 2 | Mean difference | CI low | CI high |
|---|---|---|---|---|---|---|---|
| Indel | Complex regions | F1-score | D-G | G-DV | 0.018604035714 | 0.0168715821193 | 0.020336489309 |
| Indel | Complex regions | F1-score | D-G | G-G | 0.010302960084 | 0.0098960161808 | 0.010709903987 |
| Indel | Complex regions | F1-score | G-D | G-DV | 0.003523521008 | 0.0015040720122 | 0.005542970005 |
| Indel | Complex regions | F1-score | G-D | G-G | -0.004777554622 | -0.0051838895401 | -0.004371219704 |
| Indel | Complex regions | F1-score | G-DV | G-G | -0.008301075630 | -0.0101188535565 | -0.006483297704 |
| Indel | Simple regions | Precision | D-D | D-DV | -0.000750405882 | -0.0009639653545 | -0.000536846410 |
| Indel | Simple regions | Precision | D-D | D-G | 0.001800670588 | 0.0013950818981 | 0.002206259278 |
| Indel | Simple regions | Precision | D-D | G-D | 0.000341850000 | 0.0001117368514 | 0.000571963149 |
| Indel | Simple regions | Precision | D-D | G-DV | -0.000336494118 | -0.0005645441629 | -0.000108444072 |
| Indel | Simple regions | Precision | D-D | G-G | -0.000047897059 | -0.0003054341506 | 0.000209640033 |
| Indel | Simple regions | Precision | D-DV | D-G | 0.002551076471 | 0.0020707693441 | 0.003031383597 |
| Indel | Simple regions | Precision | D-DV | G-D | 0.001092255882 | 0.0008691381161 | 0.001315373649 |
| Indel | Simple regions | Precision | D-DV | G-DV | 0.000413911765 | 0.0002810286459 | 0.000546794883 |
| Indel | Simple regions | Precision | D-DV | G-G | 0.000702508824 | 0.0005250895269 | 0.000879928120 |

| Type | Region | Metric | Pipeline 1 | Pipeline 2 | Mean difference | CI low | CI high |
|---|---|---|---|---|---|---|---|
| Indel | Simple regions | Precision | D-G | G-D | -0.001458820588 | -0.0018647552350 | -0.001052885941 |
| Indel | Simple regions | Precision | D-G | G-DV | -0.002137164706 | -0.0026027803546 | -0.001671549057 |
| Indel | Simple regions | Precision | D-G | G-G | -0.001848567647 | -0.0022838182309 | -0.001413317063 |
| Indel | Simple regions | Precision | G-D | G-DV | -0.000678344118 | -0.0009083438012 | -0.000448344434 |
| Indel | Simple regions | Precision | G-D | G-G | -0.000389747059 | -0.0006157500923 | -0.000163744025 |
| Indel | Simple regions | Precision | G-DV | G-G | 0.000288597059 | 0.0000734124555 | 0.000503781662 |
| Indel | Simple regions | Recall | D-D | D-DV | 0.001631776471 | 0.0012093944731 | 0.002054158468 |
| Indel | Simple regions | Recall | D-D | D-G | 0.000057122059 | -0.0000831107574 | 0.000197354875 |
| Indel | Simple regions | Recall | D-D | G-D | 0.011145985294 | 0.0103932640158 | 0.011898706572 |
| Indel | Simple regions | Recall | D-D | G-DV | 0.012236929412 | 0.0114573911343 | 0.013016467689 |
| Indel | Simple regions | Recall | D-D | G-G | 0.010357741176 | 0.0095921816645 | 0.011123300688 |
| Indel | Simple regions | Recall | D-DV | D-G | -0.001574654412 | -0.0019859224198 | -0.001163386404 |
| Indel | Simple regions | Recall | D-DV | G-D | 0.009514208824 | 0.0086115091175 | 0.010416908530 |
| Indel | Simple regions | Recall | D-DV | G-DV | 0.010605152941 | 0.0098405013686 | 0.011369804514 |

| Type | Region | Metric | Pipeline 1 | Pipeline 2 | Mean difference | CI low | CI high |
|---|---|---|---|---|---|---|---|
| Indel | Simple regions | Recall | D-DV | G-G | 0.008725964706 | 0.0077720365435 | 0.009679892868 |
| Indel | Simple regions | Recall | D-G | G-D | 0.011088863235 | 0.0103360730027 | 0.011841653468 |
| Indel | Simple regions | Recall | D-G | G-DV | 0.012179807353 | 0.0114031522734 | 0.012956462432 |
| Indel | Simple regions | Recall | D-G | G-G | 0.010300619118 | 0.0095444255900 | 0.011056812645 |
| Indel | Simple regions | Recall | G-D | G-DV | 0.001090944118 | 0.0007043430773 | 0.001477545158 |
| Indel | Simple regions | Recall | G-D | G-G | -0.000788244118 | -0.0009760545903 | -0.000600433645 |
| Indel | Simple regions | Recall | G-DV | G-G | -0.001879188235 | -0.0023686466152 | -0.001389729855 |
| Indel | Simple regions | F1-score | D-D | D-DV | 0.000643302521 | 0.0003467799977 | 0.000939825044 |
| Indel | Simple regions | F1-score | D-D | D-G | 0.001338096639 | 0.0010183040470 | 0.001657889230 |
| Indel | Simple regions | F1-score | D-D | G-D | 0.008278224790 | 0.0078777350874 | 0.008678714492 |
| Indel | Simple regions | F1-score | D-D | G-DV | 0.008588111345 | 0.0081697586610 | 0.009006464028 |
| Indel | Simple regions | F1-score | D-D | G-G | 0.007433250000 | 0.0069538318350 | 0.007912668165 |
| Indel | Simple regions | F1-score | D-DV | D-G | 0.000694794118 | 0.0004024286862 | 0.000987159549 |
| Indel | Simple regions | F1-score | D-DV | G-D | 0.007634922269 | 0.0071206690606 | 0.008149175477 |

| Type | Region | Metric | Pipeline 1 | Pipeline 2 | Mean difference | CI low | CI high |
|---|---|---|---|---|---|---|---|
| Indel | Simple regions | F1-score | D-DV | G-DV | 0.007944808824 | 0.0075098567323 | 0.008379760915 |
| Indel | Simple regions | F1-score | D-DV | G-G | 0.006789947479 | 0.0061814865152 | 0.007398408443 |
| Indel | Simple regions | F1-score | D-G | G-D | 0.006940128151 | 0.0064343218614 | 0.007445934441 |
| Indel | Simple regions | F1-score | D-G | G-DV | 0.007250014706 | 0.0067828174703 | 0.007717211941 |
| Indel | Simple regions | F1-score | D-G | G-G | 0.006095153361 | 0.0055118185424 | 0.006678488180 |
| Indel | Simple regions | F1-score | G-D | G-DV | 0.000309886555 | -0.0000003123001 | 0.000620085409 |
| Indel | Simple regions | F1-score | G-D | G-G | -0.000844974790 | -0.0011158756805 | -0.000574073899 |
| Indel | Simple regions | F1-score | G-DV | G-G | -0.001154861345 | -0.0015593388930 | -0.000750383796 |
| Indel | Non-coding regions | Precision | D-D | D-DV | -0.000642501471 | -0.0009182099443 | -0.000366792997 |
| Indel | Non-coding regions | Precision | D-D | D-G | 0.002167710294 | 0.0018795838515 | 0.002455836737 |
| Indel | Non-coding regions | Precision | D-D | G-D | 0.004016692647 | 0.0035978696847 | 0.004435515609 |
| Indel | Non-coding regions | Precision | D-D | G-DV | 0.000754539706 | 0.0004539035425 | 0.001055175869 |
| Indel | Non-coding regions | Precision | D-D | G-G | 0.000884355882 | 0.0005271710288 | 0.001241540736 |
| Indel | Non-coding regions | Precision | D-DV | D-G | 0.002810211765 | 0.0023991720090 | 0.003221251520 |

| Type | Region | Metric | Pipeline 1 | Pipeline 2 | Mean difference | CI low | CI high |
|---|---|---|---|---|---|---|---|
| Indel | Non-coding regions | Precision | D-DV | G-D | 0.004659194118 | 0.0043071057623 | 0.005011282473 |
| Indel | Non-coding regions | Precision | D-DV | G-DV | 0.001397041176 | 0.0011893768389 | 0.001604705514 |
| Indel | Non-coding regions | Precision | D-DV | G-G | 0.001526857353 | 0.0012835828070 | 0.001770131899 |
| Indel | Non-coding regions | Precision | D-G | G-D | 0.001848982353 | 0.0013561461480 | 0.002341818558 |
| Indel | Non-coding regions | Precision | D-G | G-DV | -0.001413170588 | -0.0018082146281 | -0.001018126548 |
| Indel | Non-coding regions | Precision | D-G | G-G | -0.001283354412 | -0.0017288624966 | -0.000837846327 |
| Indel | Non-coding regions | Precision | G-D | G-DV | -0.003262152941 | -0.0036660417550 | -0.002858264127 |
| Indel | Non-coding regions | Precision | G-D | G-G | -0.003132336765 | -0.0034072018140 | -0.002857471715 |
| Indel | Non-coding regions | Precision | G-DV | G-G | 0.000129816176 | -0.0002034787731 | 0.000463111126 |
| Indel | Non-coding regions | Recall | D-D | D-DV | 0.008777817647 | 0.0074565582499 | 0.010099077044 |
| Indel | Non-coding regions | Recall | D-D | D-G | 0.003116401471 | 0.0028135721186 | 0.003419230823 |
| Indel | Non-coding regions | Recall | D-D | G-D | 0.019201002941 | 0.0181322565018 | 0.020269749381 |
| Indel | Non-coding regions | Recall | D-D | G-DV | 0.024528923529 | 0.0225916650485 | 0.026466182010 |
| Indel | Non-coding regions | Recall | D-D | G-G | 0.016975835294 | 0.0159910467895 | 0.017960623799 |

| Type | Region | Metric | Pipeline 1 | Pipeline 2 | Mean difference | CI low | CI high |
|---|---|---|---|---|---|---|---|
| Indel | Non-coding regions | Recall | D-DV | D-G | -0.005661416176 | -0.0069712336630 | -0.004351598690 |
| Indel | Non-coding regions | Recall | D-DV | G-D | 0.010423185294 | 0.0089125924827 | 0.011933778106 |
| Indel | Non-coding regions | Recall | D-DV | G-DV | 0.015751105882 | 0.0148137909113 | 0.016688420853 |
| Indel | Non-coding regions | Recall | D-DV | G-G | 0.008198017647 | 0.0067955101903 | 0.009600525104 |
| Indel | Non-coding regions | Recall | D-G | G-D | 0.016084601471 | 0.0152094327629 | 0.016959770178 |
| Indel | Non-coding regions | Recall | D-G | G-DV | 0.021412522059 | 0.0195699866777 | 0.023255057440 |
| Indel | Non-coding regions | Recall | D-G | G-G | 0.013859433824 | 0.0130827742674 | 0.014636093380 |
| Indel | Non-coding regions | Recall | G-D | G-DV | 0.005327920588 | 0.0037330197192 | 0.006922821457 |
| Indel | Non-coding regions | Recall | G-D | G-G | -0.002225167647 | -0.0024738876756 | -0.001976447618 |
| Indel | Non-coding regions | Recall | G-DV | G-G | -0.007553088235 | -0.0090772838564 | -0.006028892614 |
| Indel | Non-coding regions | F1-score | D-D | D-DV | 0.006126983193 | 0.0052749544506 | 0.006979011936 |
| Indel | Non-coding regions | F1-score | D-D | D-G | 0.003768920168 | 0.0034855131963 | 0.004052327140 |
| Indel | Non-coding regions | F1-score | D-D | G-D | 0.016790661765 | 0.0163264805956 | 0.017254842934 |
| Indel | Non-coding regions | F1-score | D-D | G-DV | 0.018740487395 | 0.0175779538188 | 0.019903020971 |

| Type | Region | Metric | Pipeline 1 | Pipeline 2 | Mean difference | CI low | CI high |
|---|---|---|---|---|---|---|---|
| Indel | Non-coding regions | F1-score | D-D | G-G | 0.012991714286 | 0.0125672040818 | 0.013416224490 |
| Indel | Non-coding regions | F1-score | D-DV | D-G | -0.002358063025 | -0.0031713347208 | -0.001544791330 |
| Indel | Non-coding regions | F1-score | D-DV | G-D | 0.010663678571 | 0.0095263556489 | 0.011801001494 |
| Indel | Non-coding regions | F1-score | D-DV | G-DV | 0.012613504202 | 0.0121043447834 | 0.013122663620 |
| Indel | Non-coding regions | F1-score | D-DV | G-G | 0.006864731092 | 0.0058161106067 | 0.007913351578 |
| Indel | Non-coding regions | F1-score | D-G | G-D | 0.013021741597 | 0.0125420936754 | 0.013501389518 |
| Indel | Non-coding regions | F1-score | D-G | G-DV | 0.014971567227 | 0.0138719088099 | 0.016071225644 |
| Indel | Non-coding regions | F1-score | D-G | G-G | 0.009222794118 | 0.0088350600797 | 0.009610528156 |
| Indel | Non-coding regions | F1-score | G-D | G-DV | 0.001949825630 | 0.0005763781721 | 0.003323273088 |
| Indel | Non-coding regions | F1-score | G-D | G-G | -0.003798947479 | -0.0040875328205 | -0.003510362137 |
| Indel | Non-coding regions | F1-score | G-DV | G-G | -0.005748773109 | -0.0070212402479 | -0.004476305971 |
| Indel | Coding regions | Precision | D-D | D-DV | 0.010514602941 | -0.0035967486692 | 0.024625954552 |
| Indel | Coding regions | Precision | D-D | D-G | 0.046653198529 | 0.0319321887688 | 0.061374208290 |
| Indel | Coding regions | Precision | D-D | G-D | 0.009548119118 | -0.0047035661939 | 0.023799804429 |

| Type | Region | Metric | Pipeline 1 | Pipeline 2 | Mean difference | CI low | CI high |
|---|---|---|---|---|---|---|---|
| Indel | Coding regions | Precision | D-D | G-DV | -0.000407822059 | -0.0133470765661 | 0.012531432448 |
| Indel | Coding regions | Precision | D-D | G-G | 0.004527338235 | -0.0094506111176 | 0.018505287588 |
| Indel | Coding regions | Precision | D-DV | D-G | 0.036138595588 | 0.0209121441684 | 0.051365047008 |
| Indel | Coding regions | Precision | D-DV | G-D | -0.000966483824 | -0.0174958494858 | 0.015562881839 |
| Indel | Coding regions | Precision | D-DV | G-DV | -0.010922425000 | -0.0217521642935 | -0.000092685707 |
| Indel | Coding regions | Precision | D-DV | G-G | -0.005987264706 | -0.0222288670301 | 0.010254337618 |
| Indel | Coding regions | Precision | D-G | G-D | -0.037105079412 | -0.0528423460635 | -0.021367812760 |
| Indel | Coding regions | Precision | D-G | G-DV | -0.047061020588 | -0.0632211424458 | -0.030900898731 |
| Indel | Coding regions | Precision | D-G | G-G | -0.042125860294 | -0.0575200161153 | -0.026731704473 |
| Indel | Coding regions | Precision | G-D | G-DV | -0.009955941176 | -0.0249102210481 | 0.004998338695 |
| Indel | Coding regions | Precision | G-D | G-G | -0.005020780882 | -0.0079782648203 | -0.002063296944 |
| Indel | Coding regions | Precision | G-DV | G-G | 0.004935160294 | -0.0097216786718 | 0.019591999260 |
| Indel | Coding regions | Recall | D-D | D-DV | 0.018273779412 | 0.0043706390638 | 0.032176919760 |
| Indel | Coding regions | Recall | D-D | D-G | 0.038245701471 | 0.0239100562862 | 0.052581346655 |

| Type | Region | Metric | Pipeline 1 | Pipeline 2 | Mean difference | CI low | CI high |
|---|---|---|---|---|---|---|---|
| Indel | Coding regions | Recall | D-D | G-D | 0.043830920588 | 0.0293763506969 | 0.058285490480 |
| Indel | Coding regions | Recall | D-D | G-DV | 0.038997227941 | 0.0256119034651 | 0.052382552417 |
| Indel | Coding regions | Recall | D-D | G-G | 0.037078047059 | 0.0229013382889 | 0.051254755829 |
| Indel | Coding regions | Recall | D-DV | D-G | 0.019971922059 | 0.0051717969695 | 0.034772047148 |
| Indel | Coding regions | Recall | D-DV | G-D | 0.025557141176 | 0.0086248268067 | 0.042489455546 |
| Indel | Coding regions | Recall | D-DV | G-DV | 0.020723448529 | 0.0089631158371 | 0.032483781222 |
| Indel | Coding regions | Recall | D-DV | G-G | 0.018804267647 | 0.0021244334563 | 0.035484101838 |
| Indel | Coding regions | Recall | D-G | G-D | 0.005585219118 | -0.0105926730399 | 0.021763111275 |
| Indel | Coding regions | Recall | D-G | G-DV | 0.000751526471 | -0.0159791431091 | 0.017482196050 |
| Indel | Coding regions | Recall | D-G | G-G | -0.001167654412 | -0.0170042155574 | 0.014668906734 |
| Indel | Coding regions | Recall | G-D | G-DV | -0.004833692647 | -0.0195905979482 | 0.009923212654 |
| Indel | Coding regions | Recall | G-D | G-G | -0.006752873529 | -0.0097243302126 | -0.003781416846 |
| Indel | Coding regions | Recall | G-DV | G-G | -0.001919180882 | -0.0164302123399 | 0.012591850575 |
| Indel | Coding regions | F1-score | D-D | D-DV | -0.002933827273 | -0.0060777900129 | 0.000210135467 |

| Type | Region | Metric | Pipeline 1 | Pipeline 2 | Mean difference | CI low | CI high |
|---|---|---|---|---|---|---|---|
| Indel | Coding regions | F1-score | D-D | D-G | 0.009759881944 | 0.0047814165768 | 0.014738347312 |
| Indel | Coding regions | F1-score | D-D | G-D | 0.034668132883 | 0.0283075284193 | 0.041028737347 |
| Indel | Coding regions | F1-score | D-D | G-DV | 0.025394690583 | 0.0205547405170 | 0.030234640649 |
| Indel | Coding regions | F1-score | D-D | G-G | 0.025248871622 | 0.0204085046879 | 0.030089238555 |
| Indel | Coding regions | F1-score | D-DV | D-G | 0.012927589202 | 0.0079677570949 | 0.017887421309 |
| Indel | Coding regions | F1-score | D-DV | G-D | 0.038441324885 | 0.0329008854491 | 0.043981764320 |
| Indel | Coding regions | F1-score | D-DV | G-DV | 0.028480623025 | 0.0251448343902 | 0.031816411659 |
| Indel | Coding regions | F1-score | D-DV | G-G | 0.028805029954 | 0.0251528209386 | 0.032457238969 |
| Indel | Coding regions | F1-score | D-G | G-D | 0.026052172494 | 0.0184643824931 | 0.033639962495 |
| Indel | Coding regions | F1-score | D-G | G-DV | 0.016611595794 | 0.0101528716261 | 0.023070319963 |
| Indel | Coding regions | F1-score | D-G | G-G | 0.016303566434 | 0.0101664724586 | 0.022440660409 |
| Indel | Coding regions | F1-score | G-D | G-DV | -0.009221358277 | -0.0142758435588 | -0.004166872994 |
| Indel | Coding regions | F1-score | G-D | G-G | -0.009232123620 | -0.0135311850625 | -0.004933062178 |
| Indel | Coding regions | F1-score | G-DV | G-G | -0.000261979592 | -0.0027825279331 | 0.002258568749 |

| Type | Region | Metric | Pipeline 1 | Pipeline 2 | Mean difference | CI low | CI high |
|---|---|---|---|---|---|---|---|
| SNV | All regions | Precision | D-D | D-DV | -0.001023691176 | -0.0010523507074 | -0.000995031646 |
| SNV | All regions | Precision | D-D | D-G | 0.000682705882 | 0.0005866871382 | 0.000778724627 |
| SNV | All regions | Precision | D-D | G-D | 0.003464735294 | 0.0033908249456 | 0.003538645643 |
| SNV | All regions | Precision | D-D | G-DV | -0.000938955882 | -0.0009800944136 | -0.000897817351 |
| SNV | All regions | Precision | D-D | G-G | 0.005518705882 | 0.0054470384945 | 0.005590373270 |
| SNV | All regions | Precision | D-DV | D-G | 0.001706397059 | 0.0016141830487 | 0.001798611069 |
| SNV | All regions | Precision | D-DV | G-D | 0.004488426471 | 0.0044037867167 | 0.004573066224 |
| SNV | All regions | Precision | D-DV | G-DV | 0.000084735294 | 0.0000439867302 | 0.000125483858 |
| SNV | All regions | Precision | D-DV | G-G | 0.006542397059 | 0.0064679922326 | 0.006616801885 |
| SNV | All regions | Precision | D-G | G-D | 0.002782029412 | 0.0026498606484 | 0.002914198175 |
| SNV | All regions | Precision | D-G | G-DV | -0.001621661765 | -0.0017275792262 | -0.001515744303 |
| SNV | All regions | Precision | D-G | G-G | 0.004836000000 | 0.0047560401744 | 0.004915959826 |
| SNV | All regions | Precision | G-D | G-DV | -0.004403691176 | -0.0044635424505 | -0.004343839902 |
| SNV | All regions | Precision | G-D | G-G | 0.002053970588 | 0.0019441121978 | 0.002163828979 |

| Type | Region | Metric | Pipeline 1 | Pipeline 2 | Mean difference | CI low | CI high |
|---|---|---|---|---|---|---|---|
| SNV | All regions | Precision | G-DV | G-G | 0.006457661765 | 0.0063736725977 | 0.006541650932 |
| SNV | All regions | Recall | D-D | D-DV | 0.000597970588 | 0.0005719971839 | 0.000623943993 |
| SNV | All regions | Recall | D-D | D-G | 0.003345470588 | 0.0033085916853 | 0.003382349491 |
| SNV | All regions | Recall | D-D | G-D | 0.023620161765 | 0.0235645544426 | 0.023675769087 |
| SNV | All regions | Recall | D-D | G-DV | 0.021002470588 | 0.0209690979722 | 0.021035843204 |
| SNV | All regions | Recall | D-D | G-G | 0.023684161765 | 0.0236339198538 | 0.023734403676 |
| SNV | All regions | Recall | D-DV | D-G | 0.002747500000 | 0.0027011980303 | 0.002793801970 |
| SNV | All regions | Recall | D-DV | G-D | 0.023022191176 | 0.0229579093579 | 0.023086472995 |
| SNV | All regions | Recall | D-DV | G-DV | 0.020404500000 | 0.0203617307622 | 0.020447269238 |
| SNV | All regions | Recall | D-DV | G-G | 0.023086191176 | 0.0230220090541 | 0.023150373299 |
| SNV | All regions | Recall | D-G | G-D | 0.020274691176 | 0.0202286387075 | 0.020320743645 |
| SNV | All regions | Recall | D-G | G-DV | 0.017657000000 | 0.0176206397260 | 0.017693360274 |
| SNV | All regions | Recall | D-G | G-G | 0.020338691176 | 0.0202942032873 | 0.020383179066 |
| SNV | All regions | Recall | G-D | G-DV | -0.002617691176 | -0.0026586678786 | -0.002576714474 |

| Type | Region | Metric | Pipeline 1 | Pipeline 2 | Mean difference | CI low | CI high |
|---|---|---|---|---|---|---|---|
| SNV | All regions | Recall | G-D | G-G | 0.000064000000 | 0.0000225705694 | 0.000105429431 |
| SNV | All regions | Recall | G-DV | G-G | 0.002681691176 | 0.0026387330304 | 0.002724649323 |
| SNV | All regions | F1-score | D-D | D-DV | -0.000211602941 | -0.0002348969903 | -0.000188308892 |
| SNV | All regions | F1-score | D-D | D-G | 0.002016941176 | 0.0019688279649 | 0.002065054388 |
| SNV | All regions | F1-score | D-D | G-D | 0.013653426471 | 0.0135970088167 | 0.013709844124 |
| SNV | All regions | F1-score | D-D | G-DV | 0.010162073529 | 0.0101304872411 | 0.010193659818 |
| SNV | All regions | F1-score | D-D | G-G | 0.014692367647 | 0.0146528982749 | 0.014731837019 |
| SNV | All regions | F1-score | D-DV | D-G | 0.002228544118 | 0.0021810995880 | 0.002275988647 |
| SNV | All regions | F1-score | D-DV | G-D | 0.013865029412 | 0.0137980552688 | 0.013932003555 |
| SNV | All regions | F1-score | D-DV | G-DV | 0.010373676471 | 0.0103366161221 | 0.010410736819 |
| SNV | All regions | F1-score | D-DV | G-G | 0.014903970588 | 0.0148592202238 | 0.014948720953 |
| SNV | All regions | F1-score | D-G | G-D | 0.011636485294 | 0.0115642126063 | 0.011708757982 |
| SNV | All regions | F1-score | D-G | G-DV | 0.008145132353 | 0.0080894848773 | 0.008200779829 |
| SNV | All regions | F1-score | D-G | G-G | 0.012675426471 | 0.0126314220566 | 0.012719430885 |

| Type | Region | Metric | Pipeline 1 | Pipeline 2 | Mean difference | CI low | CI high |
|---|---|---|---|---|---|---|---|
| SNV | All regions | F1-score | G-D | G-DV | -0.003491352941 | -0.0035339503582 | -0.003448755524 |
| SNV | All regions | F1-score | G-D | G-G | 0.001038941176 | 0.0009827237389 | 0.001095158614 |
| SNV | All regions | F1-score | G-DV | G-G | 0.004530294118 | 0.0044878783694 | 0.004572709866 |
| SNV | Complex regions | Precision | D-D | D-DV | -0.003816397059 | -0.0039212655816 | -0.003711528536 |
| SNV | Complex regions | Precision | D-D | D-G | 0.000053852941 | -0.0001679960229 | 0.000275701905 |
| SNV | Complex regions | Precision | D-D | G-D | 0.013088955882 | 0.0128257051375 | 0.013352206627 |
| SNV | Complex regions | Precision | D-D | G-DV | -0.003788705882 | -0.0039479601816 | -0.003629451583 |
| SNV | Complex regions | Precision | D-D | G-G | 0.020281544118 | 0.0200499413800 | 0.020513146855 |
| SNV | Complex regions | Precision | D-DV | D-G | 0.003870250000 | 0.0036699946944 | 0.004070505306 |
| SNV | Complex regions | Precision | D-DV | G-D | 0.016905352941 | 0.0166020228753 | 0.017208683007 |
| SNV | Complex regions | Precision | D-DV | G-DV | 0.000027691176 | -0.0001341196468 | 0.000189502000 |
| SNV | Complex regions | Precision | D-DV | G-G | 0.024097941176 | 0.0238618583419 | 0.024334024011 |
| SNV | Complex regions | Precision | D-G | G-D | 0.013035102941 | 0.0126486117665 | 0.013421594116 |
| SNV | Complex regions | Precision | D-G | G-DV | -0.003842558824 | -0.0041192679096 | -0.003565849737 |

| Type | Region | Metric | Pipeline 1 | Pipeline 2 | Mean difference | CI low | CI high |
|---|---|---|---|---|---|---|---|
| SNV | Complex regions | Precision | D-G | G-G | 0.020227691176 | 0.0199746138692 | 0.020480768484 |
| SNV | Complex regions | Precision | G-D | G-DV | -0.016877661765 | -0.0170807186178 | -0.016674604912 |
| SNV | Complex regions | Precision | G-D | G-G | 0.007192588235 | 0.0068216707169 | 0.007563505754 |
| SNV | Complex regions | Precision | G-DV | G-G | 0.024070250000 | 0.0237915647982 | 0.024348935202 |
| SNV | Complex regions | Recall | D-D | D-DV | 0.002298029412 | 0.0021950839549 | 0.002400974869 |
| SNV | Complex regions | Recall | D-D | D-G | 0.012671176471 | 0.0125283062778 | 0.012814046663 |
| SNV | Complex regions | Recall | D-D | G-D | 0.047312838235 | 0.0471228952347 | 0.047502781236 |
| SNV | Complex regions | Recall | D-D | G-DV | 0.037938352941 | 0.0378179311002 | 0.038058774782 |
| SNV | Complex regions | Recall | D-D | G-G | 0.047766676471 | 0.0475781209391 | 0.047955232002 |
| SNV | Complex regions | Recall | D-DV | D-G | 0.010373147059 | 0.0101942169504 | 0.010552077167 |
| SNV | Complex regions | Recall | D-DV | G-D | 0.045014808824 | 0.0447885000993 | 0.045241117548 |
| SNV | Complex regions | Recall | D-DV | G-DV | 0.035640323529 | 0.0354844289896 | 0.035796218069 |
| SNV | Complex regions | Recall | D-DV | G-G | 0.045468647059 | 0.0452278877949 | 0.045709406323 |
| SNV | Complex regions | Recall | D-G | G-D | 0.034641661765 | 0.0344801630264 | 0.034803160503 |

| Type | Region | Metric | Pipeline 1 | Pipeline 2 | Mean difference | CI low | CI high |
|---|---|---|---|---|---|---|---|
| SNV | Complex regions | Recall | D-G | G-DV | 0.025267176471 | 0.0251310533696 | 0.025403299572 |
| SNV | Complex regions | Recall | D-G | G-G | 0.035095500000 | 0.0349243367704 | 0.035266663230 |
| SNV | Complex regions | Recall | G-D | G-DV | -0.009374485294 | -0.0095107569124 | -0.009238213676 |
| SNV | Complex regions | Recall | G-D | G-G | 0.000453838235 | 0.0003059622597 | 0.000601714211 |
| SNV | Complex regions | Recall | G-DV | G-G | 0.009828323529 | 0.0096637073339 | 0.009992939725 |
| SNV | Complex regions | F1-score | D-D | D-DV | -0.000740411765 | -0.0008291988914 | -0.000651624638 |
| SNV | Complex regions | F1-score | D-D | D-G | 0.006422661765 | 0.0063029292572 | 0.006542394272 |
| SNV | Complex regions | F1-score | D-D | G-D | 0.030560735294 | 0.0303672118932 | 0.030754258695 |
| SNV | Complex regions | F1-score | D-D | G-DV | 0.017587897059 | 0.0174680695379 | 0.017707724580 |
| SNV | Complex regions | F1-score | D-D | G-G | 0.034265823529 | 0.0341311848079 | 0.034400462251 |
| SNV | Complex regions | F1-score | D-DV | D-G | 0.007163073529 | 0.0070390617951 | 0.007287085264 |
| SNV | Complex regions | F1-score | D-DV | G-D | 0.031301147059 | 0.0310651212869 | 0.031537172831 |
| SNV | Complex regions | F1-score | D-DV | G-DV | 0.018328308824 | 0.0181871267567 | 0.018469490890 |
| SNV | Complex regions | F1-score | D-DV | G-G | 0.035006235294 | 0.0348538397208 | 0.035158630867 |

| Type | Region | Metric | Pipeline 1 | Pipeline 2 | Mean difference | CI low | CI high |
|---|---|---|---|---|---|---|---|
| SNV | Complex regions | F1-score | D-G | G-D | 0.024138073529 | 0.0239239711421 | 0.024352175917 |
| SNV | Complex regions | F1-score | D-G | G-DV | 0.011165235294 | 0.0110172696943 | 0.011313200894 |
| SNV | Complex regions | F1-score | D-G | G-G | 0.027843161765 | 0.0276974319796 | 0.027988891550 |
| SNV | Complex regions | F1-score | G-D | G-DV | -0.012972838235 | -0.0131087000484 | -0.012836976422 |
| SNV | Complex regions | F1-score | G-D | G-G | 0.003705088235 | 0.0035262470981 | 0.003883929372 |
| SNV | Complex regions | F1-score | G-DV | G-G | 0.016677926471 | 0.0165425071235 | 0.016813345818 |
| SNV | Simple regions | Precision | D-D | D-DV | -0.000038000000 | -0.0000446021540 | -0.000031397846 |
| SNV | Simple regions | Precision | D-D | D-G | 0.000935102941 | 0.0008640249127 | 0.001006180970 |
| SNV | Simple regions | Precision | D-D | G-D | 0.000217647059 | 0.0001940383847 | 0.000241255733 |
| SNV | Simple regions | Precision | D-D | G-DV | 0.000100588235 | 0.0000922113327 | 0.000108965138 |
| SNV | Simple regions | Precision | D-D | G-G | 0.000459808824 | 0.0004134860695 | 0.000506131578 |
| SNV | Simple regions | Precision | D-DV | D-G | 0.000973102941 | 0.0009016053078 | 0.001044600575 |
| SNV | Simple regions | Precision | D-DV | G-D | 0.000255647059 | 0.0002316564084 | 0.000279637709 |
| SNV | Simple regions | Precision | D-DV | G-DV | 0.000138588235 | 0.0001323619149 | 0.000144814556 |

| Type | Region | Metric | Pipeline 1 | Pipeline 2 | Mean difference | CI low | CI high |
|---|---|---|---|---|---|---|---|
| SNV | Simple regions | Precision | D-DV | G-G | 0.000497808824 | 0.0004503504020 | 0.000545267245 |
| SNV | Simple regions | Precision | D-G | G-D | -0.000717455882 | -0.0007880304211 | -0.000646881344 |
| SNV | Simple regions | Precision | D-G | G-DV | -0.000834514706 | -0.0009044585950 | -0.000764570817 |
| SNV | Simple regions | Precision | D-G | G-G | -0.000475294118 | -0.0005189978872 | -0.000431590348 |
| SNV | Simple regions | Precision | G-D | G-DV | -0.000117058824 | -0.0001408153466 | -0.000093302300 |
| SNV | Simple regions | Precision | G-D | G-G | 0.000242161765 | 0.0001948567218 | 0.000289466808 |
| SNV | Simple regions | Precision | G-DV | G-G | 0.000359220588 | 0.0003134380406 | 0.000405003136 |
| SNV | Simple regions | Recall | D-D | D-DV | 0.000001926471 | -0.0000020049155 | 0.000005857857 |
| SNV | Simple regions | Recall | D-D | D-G | 0.000076661765 | 0.0000699551895 | 0.000083368340 |
| SNV | Simple regions | Recall | D-D | G-D | 0.015315470588 | 0.0152944034676 | 0.015336537709 |
| SNV | Simple regions | Recall | D-D | G-DV | 0.015066073529 | 0.0150560672082 | 0.015076079851 |
| SNV | Simple regions | Recall | D-D | G-G | 0.015242852941 | 0.0152308697182 | 0.015254836164 |
| SNV | Simple regions | Recall | D-DV | D-G | 0.000074735294 | 0.0000681829150 | 0.000081287673 |
| SNV | Simple regions | Recall | D-DV | G-D | 0.015313544118 | 0.0152919383402 | 0.015335149895 |

| Type | Region | Metric | Pipeline 1 | Pipeline 2 | Mean difference | CI low | CI high |
|---|---|---|---|---|---|---|---|
| SNV | Simple regions | Recall | D-DV | G-DV | 0.015064147059 | 0.0150551615400 | 0.015073132578 |
| SNV | Simple regions | Recall | D-DV | G-G | 0.015240926471 | 0.0152294619294 | 0.015252391012 |
| SNV | Simple regions | Recall | D-G | G-D | 0.015238808824 | 0.0152169762962 | 0.015260641351 |
| SNV | Simple regions | Recall | D-G | G-DV | 0.014989411765 | 0.0149779006438 | 0.015000922886 |
| SNV | Simple regions | Recall | D-G | G-G | 0.015166191176 | 0.0151538136681 | 0.015178568685 |
| SNV | Simple regions | Recall | G-D | G-DV | -0.000249397059 | -0.0002721275324 | -0.000226666585 |
| SNV | Simple regions | Recall | G-D | G-G | -0.000072617647 | -0.0000939111238 | -0.000051324170 |
| SNV | Simple regions | Recall | G-DV | G-G | 0.000176779412 | 0.0001656125064 | 0.000187946317 |
| SNV | Simple regions | F1-score | D-D | D-DV | -0.000018058824 | -0.0000219273295 | -0.000014190318 |
| SNV | Simple regions | F1-score | D-D | D-G | 0.000506029412 | 0.0004706025443 | 0.000541456279 |
| SNV | Simple regions | F1-score | D-D | G-D | 0.007823691176 | 0.0078033108500 | 0.007844071503 |
| SNV | Simple regions | F1-score | D-D | G-DV | 0.007639411765 | 0.0076331522471 | 0.007645671282 |
| SNV | Simple regions | F1-score | D-D | G-G | 0.007906058824 | 0.0078809144525 | 0.007931203195 |
| SNV | Simple regions | F1-score | D-DV | D-G | 0.000524088235 | 0.0004885338163 | 0.000559642654 |

| Type | Region | Metric | Pipeline 1 | Pipeline 2 | Mean difference | CI low | CI high |
|---|---|---|---|---|---|---|---|
| SNV | Simple regions | F1-score | D-DV | G-D | 0.007841750000 | 0.0078209808204 | 0.007862519180 |
| SNV | Simple regions | F1-score | D-DV | G-DV | 0.007657470588 | 0.0076522076484 | 0.007662733528 |
| SNV | Simple regions | F1-score | D-DV | G-G | 0.007924117647 | 0.0078986764052 | 0.007949558889 |
| SNV | Simple regions | F1-score | D-G | G-D | 0.007317661765 | 0.0072777039468 | 0.007357619583 |
| SNV | Simple regions | F1-score | D-G | G-DV | 0.007133382353 | 0.0070971499119 | 0.007169614794 |
| SNV | Simple regions | F1-score | D-G | G-G | 0.007400029412 | 0.0073765491632 | 0.007423509660 |
| SNV | Simple regions | F1-score | G-D | G-DV | -0.000184279412 | -0.0002061657279 | -0.000162393096 |
| SNV | Simple regions | F1-score | G-D | G-G | 0.000082367647 | 0.0000516753413 | 0.000113059953 |
| SNV | Simple regions | F1-score | G-DV | G-G | 0.000266647059 | 0.0002413222176 | 0.000291971900 |
| SNV | Non-coding regions | Precision | D-D | D-DV | -0.001028794118 | -0.0010575102324 | -0.001000078003 |
| SNV | Non-coding regions | Precision | D-D | D-G | 0.000671588235 | 0.0005762799662 | 0.000766896504 |
| SNV | Non-coding regions | Precision | D-D | G-D | 0.003468102941 | 0.0033944757215 | 0.003541730161 |
| SNV | Non-coding regions | Precision | D-D | G-DV | -0.000954338235 | -0.0009952418313 | -0.000913434639 |
| SNV | Non-coding regions | Precision | D-D | G-G | 0.005498088235 | 0.0054257831978 | 0.005570393273 |

| Type | Region | Metric | Pipeline 1 | Pipeline 2 | Mean difference | CI low | CI high |
|---|---|---|---|---|---|---|---|
| SNV | Non-coding regions | Precision | D-DV | D-G | 0.001700382353 | 0.0016089755581 | 0.001791789148 |
| SNV | Non-coding regions | Precision | D-DV | G-D | 0.004496897059 | 0.0044119607624 | 0.004581833355 |
| SNV | Non-coding regions | Precision | D-DV | G-DV | 0.000074455882 | 0.0000337073438 | 0.000115204421 |
| SNV | Non-coding regions | Precision | D-DV | G-G | 0.006526882353 | 0.0064517006190 | 0.006602064087 |
| SNV | Non-coding regions | Precision | D-G | G-D | 0.002796514706 | 0.0026645100513 | 0.002928519360 |
| SNV | Non-coding regions | Precision | D-G | G-DV | -0.001625926471 | -0.0017314536478 | -0.001520399293 |
| SNV | Non-coding regions | Precision | D-G | G-G | 0.004826500000 | 0.0047446245739 | 0.004908375426 |
| SNV | Non-coding regions | Precision | G-D | G-DV | -0.004422441176 | -0.0044831160820 | -0.004361766271 |
| SNV | Non-coding regions | Precision | G-D | G-G | 0.002029985294 | 0.0019189222460 | 0.002141048342 |
| SNV | Non-coding regions | Precision | G-DV | G-G | 0.006452426471 | 0.0063679483640 | 0.006536904577 |
| SNV | Non-coding regions | Recall | D-D | D-DV | 0.000597852941 | 0.0005714514364 | 0.000624254446 |
| SNV | Non-coding regions | Recall | D-D | D-G | 0.003328426471 | 0.0032914371688 | 0.003365415772 |
| SNV | Non-coding regions | Recall | D-D | G-D | 0.023471691176 | 0.0234158468446 | 0.023527535508 |
| SNV | Non-coding regions | Recall | D-D | G-DV | 0.020846779412 | 0.0208133992141 | 0.020880159609 |

| Type | Region | Metric | Pipeline 1 | Pipeline 2 | Mean difference | CI low | CI high |
|---|---|---|---|---|---|---|---|
| SNV | Non-coding regions | Recall | D-D | G-G | 0.023543455882 | 0.0234937734062 | 0.023593138359 |
| SNV | Non-coding regions | Recall | D-DV | D-G | 0.002730573529 | 0.0026846139112 | 0.002776533148 |
| SNV | Non-coding regions | Recall | D-DV | G-D | 0.022873838235 | 0.0228100096756 | 0.022937666795 |
| SNV | Non-coding regions | Recall | D-DV | G-DV | 0.020248926471 | 0.0202068863004 | 0.020290966641 |
| SNV | Non-coding regions | Recall | D-DV | G-G | 0.022945602941 | 0.0228822335429 | 0.023008972339 |
| SNV | Non-coding regions | Recall | D-G | G-D | 0.020143264706 | 0.0200961932134 | 0.020190336198 |
| SNV | Non-coding regions | Recall | D-G | G-DV | 0.017518352941 | 0.0174816686488 | 0.017555037234 |
| SNV | Non-coding regions | Recall | D-G | G-G | 0.020215029412 | 0.0201700718658 | 0.020259986958 |
| SNV | Non-coding regions | Recall | G-D | G-DV | -0.002624911765 | -0.0026660927510 | -0.002583730778 |
| SNV | Non-coding regions | Recall | G-D | G-G | 0.000071764706 | 0.0000302916249 | 0.000113237787 |
| SNV | Non-coding regions | Recall | G-DV | G-G | 0.002696676471 | 0.0026536754518 | 0.002739677489 |
| SNV | Non-coding regions | F1-score | D-D | D-DV | -0.000214147059 | -0.0002376200177 | -0.000190674100 |
| SNV | Non-coding regions | F1-score | D-D | D-G | 0.002002808824 | 0.0019547778815 | 0.002050839766 |
| SNV | Non-coding regions | F1-score | D-D | G-D | 0.013579147059 | 0.0135228680047 | 0.013635426113 |

| Type | Region | Metric | Pipeline 1 | Pipeline 2 | Mean difference | CI low | CI high |
|---|---|---|---|---|---|---|---|
| SNV | Non-coding regions | F1-score | D-D | G-DV | 0.010074735294 | 0.0100433335858 | 0.010106137002 |
| SNV | Non-coding regions | F1-score | D-D | G-G | 0.014610455882 | 0.0145711240132 | 0.014649787751 |
| SNV | Non-coding regions | F1-score | D-DV | D-G | 0.002216955882 | 0.0021699023982 | 0.002264009367 |
| SNV | Non-coding regions | F1-score | D-DV | G-D | 0.013793294118 | 0.0137265616454 | 0.013860026590 |
| SNV | Non-coding regions | F1-score | D-DV | G-DV | 0.010288882353 | 0.0102523604492 | 0.010325404257 |
| SNV | Non-coding regions | F1-score | D-DV | G-G | 0.014824602941 | 0.0147803167911 | 0.014868889091 |
| SNV | Non-coding regions | F1-score | D-G | G-D | 0.011576338235 | 0.0115038050057 | 0.011648871465 |
| SNV | Non-coding regions | F1-score | D-G | G-DV | 0.008071926471 | 0.0080162824772 | 0.008127570464 |
| SNV | Non-coding regions | F1-score | D-G | G-G | 0.012607647059 | 0.0125628023177 | 0.012652491800 |
| SNV | Non-coding regions | F1-score | G-D | G-DV | -0.003504411765 | -0.0035474951826 | -0.003461328347 |
| SNV | Non-coding regions | F1-score | G-D | G-G | 0.001031308824 | 0.0009744738014 | 0.001088143846 |
| SNV | Non-coding regions | F1-score | G-DV | G-G | 0.004535720588 | 0.0044933822266 | 0.004578058950 |
| SNV | Coding regions | Precision | D-D | D-DV | -0.000403426471 | -0.0005908477869 | -0.000216005154 |
| SNV | Coding regions | Precision | D-D | D-G | 0.002035352941 | 0.0014632074433 | 0.002607498439 |

| Type | Region | Metric | Pipeline 1 | Pipeline 2 | Mean difference | CI low | CI high |
|---|---|---|---|---|---|---|---|
| SNV | Coding regions | Precision | D-D | G-D | 0.003023588235 | 0.0025966609456 | 0.003450515525 |
| SNV | Coding regions | Precision | D-D | G-DV | 0.000976088235 | 0.0006429392766 | 0.001309237194 |
| SNV | Coding regions | Precision | D-D | G-G | 0.008070808824 | 0.0074478875455 | 0.008693730102 |
| SNV | Coding regions | Precision | D-DV | D-G | 0.002438779412 | 0.0018568994181 | 0.003020659405 |
| SNV | Coding regions | Precision | D-DV | G-D | 0.003427014706 | 0.0030608909021 | 0.003793138510 |
| SNV | Coding regions | Precision | D-DV | G-DV | 0.001379514706 | 0.0011020869292 | 0.001656942483 |
| SNV | Coding regions | Precision | D-DV | G-G | 0.008474235294 | 0.0078691260707 | 0.009079344518 |
| SNV | Coding regions | Precision | D-G | G-D | 0.000988235294 | 0.0003019611143 | 0.001674509474 |
| SNV | Coding regions | Precision | D-G | G-DV | -0.001059264706 | -0.0017160139615 | -0.000402515450 |
| SNV | Coding regions | Precision | D-G | G-G | 0.006035455882 | 0.0052041788069 | 0.006866732958 |
| SNV | Coding regions | Precision | G-D | G-DV | -0.002047500000 | -0.0024565094293 | -0.001638490571 |
| SNV | Coding regions | Precision | G-D | G-G | 0.005047220588 | 0.0044288615675 | 0.005665579609 |
| SNV | Coding regions | Precision | G-DV | G-G | 0.007094720588 | 0.0065197895783 | 0.007669651598 |
| SNV | Coding regions | Recall | D-D | D-DV | 0.000610279412 | 0.0003425576763 | 0.000878001147 |

| Type | Region | Metric | Pipeline 1 | Pipeline 2 | Mean difference | CI low | CI high |
|---|---|---|---|---|---|---|---|
| SNV | Coding regions | Recall | D-D | D-G | 0.005423485294 | 0.0050049942828 | 0.005841976305 |
| SNV | Coding regions | Recall | D-D | G-D | 0.041779367647 | 0.0413786347031 | 0.042180100591 |
| SNV | Coding regions | Recall | D-D | G-DV | 0.040040632353 | 0.0397020650056 | 0.040379199700 |
| SNV | Coding regions | Recall | D-D | G-G | 0.040892735294 | 0.0404560422657 | 0.041329428323 |
| SNV | Coding regions | Recall | D-DV | D-G | 0.004813205882 | 0.0043579528749 | 0.005268458890 |
| SNV | Coding regions | Recall | D-DV | G-D | 0.041169088235 | 0.0407281570917 | 0.041610019379 |
| SNV | Coding regions | Recall | D-DV | G-DV | 0.039430352941 | 0.0391016515989 | 0.039759054283 |
| SNV | Coding regions | Recall | D-DV | G-G | 0.040282455882 | 0.0398270866857 | 0.040737825079 |
| SNV | Coding regions | Recall | D-G | G-D | 0.036355882353 | 0.0358855194146 | 0.036826245291 |
| SNV | Coding regions | Recall | D-G | G-DV | 0.034617147059 | 0.0341650972663 | 0.035069196851 |
| SNV | Coding regions | Recall | D-G | G-G | 0.035469250000 | 0.0349812682165 | 0.035957231783 |
| SNV | Coding regions | Recall | G-D | G-DV | -0.001738735294 | -0.0021001957273 | -0.001377274861 |
| SNV | Coding regions | Recall | G-D | G-G | -0.000886632353 | -0.0012005650581 | -0.000572699648 |
| SNV | Coding regions | Recall | G-DV | G-G | 0.000852102941 | 0.0004748342821 | 0.001229371600 |

| Type | Region | Metric | Pipeline 1 | Pipeline 2 | Mean difference | CI low | CI high |
|---|---|---|---|---|---|---|---|
| SNV | Coding regions | F1-score | D-D | D-DV | 0.000105455882 | -0.0000880498174 | 0.000298961582 |
| SNV | Coding regions | F1-score | D-D | D-G | 0.003740220588 | 0.0033777642789 | 0.004102676898 |
| SNV | Coding regions | F1-score | D-D | G-D | 0.022855676471 | 0.0225262601835 | 0.023185092758 |
| SNV | Coding regions | F1-score | D-D | G-DV | 0.020967735294 | 0.0207045421115 | 0.021230928477 |
| SNV | Coding regions | F1-score | D-D | G-G | 0.024817779412 | 0.0243913968717 | 0.025244161952 |
| SNV | Coding regions | F1-score | D-DV | D-G | 0.003634764706 | 0.0032620410930 | 0.004007488319 |
| SNV | Coding regions | F1-score | D-DV | G-D | 0.022750220588 | 0.0224445830715 | 0.023055858105 |
| SNV | Coding regions | F1-score | D-DV | G-DV | 0.020862279412 | 0.0206137121479 | 0.021110846676 |
| SNV | Coding regions | F1-score | D-DV | G-G | 0.024712323529 | 0.0242980148511 | 0.025126632208 |
| SNV | Coding regions | F1-score | D-G | G-D | 0.019115455882 | 0.0187056505661 | 0.019525261199 |
| SNV | Coding regions | F1-score | D-G | G-DV | 0.017227514706 | 0.0168577374714 | 0.017597291940 |
| SNV | Coding regions | F1-score | D-G | G-G | 0.021077558824 | 0.0205778145329 | 0.021577303114 |
| SNV | Coding regions | F1-score | G-D | G-DV | -0.001887941176 | -0.0021789706216 | -0.001596911731 |
| SNV | Coding regions | F1-score | G-D | G-G | 0.001962102941 | 0.0015829344020 | 0.002341271480 |

| Type | Region | Metric | Pipeline 1 | Pipeline 2 | Mean difference | CI low | CI high |
|---|---|---|---|---|---|---|---|
| SNV | Coding regions | F1-score | G-DV | G-G | 0.003850044118 | 0.0034730736802 | 0.004227014555 |
